# Mitochondrial and redox modifications in early stages of Huntington’s disease

**DOI:** 10.1101/2022.01.14.476381

**Authors:** Carla Lopes, I. Luísa Ferreira, Carina Maranga, Margarida Beatriz, Sandra I. Mota, José Sereno, João Castelhano, Antero Abrunhosa, Francisco Oliveira, Maura De Rosa, Michael Hayden, Mário N. Laço, Cristina Januário, Miguel Castelo Branco, A. Cristina Rego

## Abstract

Defects in mitochondrial function and mitochondrial-related redox deregulation have been attributed to Huntington’s disease (HD), a genetic neurodegenerative disorder largely affecting the striatum. However, whether these changes occur in early stages of the disease and can be detected *in vivo* is still unclear. Thus, in the present study, we analyzed changes in mitochondrial function and overreduced states associated with production of reactive oxygen species (ROS) at early stages and along disease progression. Studies were performed *in vivo* in human brain using positron emission tomography (PET) using [^64^Cu]-ATSM and *ex vivo* in human skin fibroblasts of premanifest and prodromal (Pre-M) and manifest HD patients; *in vivo* brain [^64^Cu]-ATSM PET and isolated mitochondria derived from striatum and cortex were also analyzed in YAC128 transgenic mouse at pre-symptomatic (3 month-old, mo) and symptomatic (6 to 12 mo) stages. Oxygen consumption rates were assessed by Seahorse analysis, hydrogen peroxide levels were determined using fluorescent probes and mitochondrial morphology by transmission electron microscopy in human skin fibroblasts and mouse striatal and cortical isolated mitochondria. Pre-M HD carriers exhibited enhanced whole-brain (with exception of caudate) [^64^Cu]-ATSM labelling, correlating with CAG repeat number. Fibroblasts from Pre-M showed enhanced basal and maximal respiration, proton (H^+^) leak and increased hydrogen peroxide levels, the later progressing to manifest HD; mitochondria from fibroblasts of Pre-M HD carriers also showed reduced roundness, while higher number of mitochondrial DNA copies correlated with maximal respiratory capacity. *In vivo* animal PET analysis showed increased accumulation of [^64^Cu]-ATSM in YAC128 mouse striatum. Pre-symptomatic YAC128 mouse striatal isolated mitochondria exhibited a rise in basal and maximal mitochondrial respiration and in ATP production, along with increased complex II and III activities; mouse HD mitochondria also showed enhanced mitochondrial hydrogen peroxide levels and roundness, as revealed by brain ultrastructure analysis, and defects in Ca^2+^ handling, supporting increased striatal susceptibility in YAC128 mouse brain. Data demonstrate both human and mouse mitochondrial overactivity and altered morphology at early HD stages, facilitating redox unbalance, the latter extending over manifest disease stages.

## INTRODUCTION

Huntington’s disease (HD) is an inherited neurodegenerative disorder characterized by psychiatric, motor and cognitive symptoms, strongly affecting the striatum and the cortex. HD is linked to a dynamic mutation, an expansion in CAG repeats, located in the exon 1 of the *HTT* gene which encodes for the huntingtin (HTT) protein ^1^. Expression of mutant HTT (mHTT) has been associated with mitochondrial dysfunction, including reduced mitochondrial transmembrane potential, bioenergetic abnormalities, defects in Ca^2+^ handling, alterations in organelle morphology and neurite movement, and increased production of reactive oxygen species (ROS)^2–4^ largely attributed to HD late symptomatic stages.

Evidence of mitochondrial bioenergetic dysfunction emerged from studies in post-mortem HD striata demonstrating defects in activities of respiratory complexes (Cx) II, III and IV and aconitase^5–7^. HD genetic models also showed mitochondrial defects, namely reduced levels and assembly of Cx II subunits, as observed in R6/1 and Htt171-82Q transgenic mice^8^. These studies supported that mitochondrial dysfunction could contribute to the hypometabolism and progressive atrophy of the caudate in HD. However, no differences were observed in respiratory activity in YAC128 mouse brain synaptic and non-synaptic mitochondria and striatal neurons^9,10^, or in isolated brain mitochondria and striatal neurons from R6/2 mice^11^. Conversely, we recently showed reduced respiratory profile and decreased mitochondrial membrane potential in YAC128 mouse cortico- and striatal neurons, and unexpected increased complexes activity in isolated striatal mitochondria from young YAC128 mouse, at 3 mo^12^. In addition, we found reduced activity of complex I and citrate synthase in mitochondrial platelets from pre-manifest HD carriers^13^. Results in skin fibroblasts from HD patients also demonstrated a reduction in ATP levels, suggesting a deficit of mitochondrial oxidative metabolism^14,15^. Gardiner and co-authors established a correlation between the age of onset and bioenergetic profile, as patients with earlier onset displayed more severe mitochondrial defects, independently of CAG repeat number^14^. These data implicate imbalanced mitochondrial activity in HD human peripheral cells.

Mitochondrial dysfunction and altered mitochondrial calcium accumulation are also interconnected. Previously, we observed increased Ca^2+^ loading capacity in brain mitochondria isolated from R6/2 and YAC128 mice^16^, and reduced Ca^2+^ handling in YAC128 mouse striatal neurons following elevated cytosolic Ca^2+^ due to selective activation of *N*-methyl-D-aspartate receptors^17^. A close relationship between mitochondrial deregulation and excessive ROS levels, namely superoxide anion and hydrogen peroxide, was also previously demonstrated in HD human cybrids (retaining HD platelet mitochondria) following a stress stimulus^18,19^. Evidences of redox deregulation were also found by us in ST*Hdh*^Q111/Q111^ striatal cells^20,21^. Other studies in HD human skin fibroblasts and lymphoblasts showed increased ROS levels and upregulation of antioxidant enzymes (e.g. superoxide dismutase 2, glutathione reductase, catalase)^15,22,23^.

Despite several findings indicating deregulated mitochondrial function in HD brain and peripheral cells, data is not totally consistent while *in vivo* mitochondrial related oxidative stress namely in living HD patients requires further elucidation. The PET radioisotope [^64^Cu]-ATSM ((64)Cu-labeled diacetyl-bis(N(4)-methylthiosemicarbazone)) has been applied to visualize regional oxidative stress in patients with mitochondrial dysfunction-associated diseases^24–26^. [^64^Cu]-ATSM accumulation occurs in electron rich environments, caused by mitochondrial impairment, as demonstrated in cells that are in overreduced states due to impaired mitochondrial electron transport chain (ETC)^27^. Therefore, to further elucidate the occurrence of mitochondrial dysfunction in HD, in the present study we investigated redox changes linked to mitochondrial deregulation both in premanifest/prodromal *versus* manifest HD patients and in YAC128 transgenic mouse model (at pre-/early-symptomatic to symptomatic stages) and its relationship with disease severity using [^64^Cu]-ATSM. Furthermore, we correlated *in vivo* brain data with analysis of mitochondrial function, production of mitochondrial hydrogen peroxide and mitochondrial morphology in human fibroblasts and in isolated mitochondria derived from YAC128 mouse brain striatum and cortex at different stages of the disease to assess their relevance during disease progression.

## MATERIALS AND METHODS

### Human study

#### Participant’s characterization

Patients and controls were selected according to clinical evaluation performed at Neurology consultation of Centro Hospitalar e Universitário de Coimbra (CHUC). Nine subjects were selected, six were mutant *HTT* gene carriers (HD carriers) with ages ranging from 25-66 years-old, including two premanifest (pre-M1 and pre-M2), one prodromal (pre-M3) and three manifest (HD1-3), as defined by an expert neurologist, based on motor, psychiatric, and cognitive symptoms, and three noncarriers/controls^28^. The age at onset was based on the date of clinical diagnosis, while the age at symptom onset was estimated on the information provided by the patient and the patient’s family (**Table 1**). This study was performed following an informed consent of all participants, by following the guidelines of the CHUC and after approval by the Ethic Committee of the Faculty of Medicine of the University of Coimbra (ref. CE-137/2015).

**TABLE 1:** Clinical data of Huntington’s disease carriers and patient’s and control groups. Abbreviations: F- female; M- male; n.a.- not applied; C- controls; pre-M- premanifest HD carriers; HD- Manifest HD patients.

|  | Code | CAG repeat number | Sex | Calendar age, y | Disease duration, y | Age at clinical diagnosis, y | Age at symptom onset, y | Symptom at onset | Symptoms UHDR |
| --- | --- | --- | --- | --- | --- | --- | --- | --- | --- |
| Control | C1 | 23 | F | 25 | n.a. | n.a. | n.a. | n.a. | n.a. |
|  | C2 | 25 | M | 46 | n.a. | n.a. | n.a. | n.a. | n.a. |
|  | C3 | 23 | M | 47 | n.a. | n.a. | n.a. | n.a. | n.a. |
| Premanifest & Prodromal HD | Pre-M1 | 46 | M | 25 | n.a. | 22 | n.a. | n.a. | 2 |
|  | Pre-M2 | 42 | M | 30 | n.a. | 29 | n.a. | n.a. | n.a. |
|  | Pre-M3 | 52 | F | 29 | 7 | 28 | 22 | cognitive | 24 |
| Manifest HD | HD1 | 43 | M | 62 | 22 | 47 | 40 | psychiatric | 30 |
|  | HD2 | 42 | F | 66 | 19 | 54 | 47 | psychiatric | Motor: 33 |
|  | HD3 | 46 | F | 43 | 7 | 39 | 36 | psychiatric | 72 |

#### PET [^64^Cu]-ATSM PET acquisition and processing

Preparation of [^64^Cu]-ATSM was performed as described elsewhere^29^. Cyclotron produced [^64^Cu]CuCl_2_ solution was added to a reactor containing H2-ATSM in CH_3_COONa 1 M. The mixture was purified in a Sep-Pak tC18 light. [^64^Cu]-ATSM was obtained in a 10% ethanol saline sterile solution, with ≥ 99% of radiochemical purity and specific activity > 200 GBq/μmol. A Philips Gemini GXL PET/CT scanner (Philips Medical Systems, Best, the Netherlands) was used to perform a dynamic 3-dimensional PET [^64^Cu]-ATSM scan of the entire brain (90 slices, 2-mm slice sampling) and a low-dose brain CT scan, for attenuation correction. Antioxidant vitamin supplements were suspended 24 hours prior to the scan. To prevent possible head movement during acquisition, patients’ head was restrained with a soft elastic tape. PET scan was acquired in a group of seven subjects (two control participants and five HD carriers) over a period of 60 minutes (24 frames: 4×15s + 4×30s + 3×60s + 2×120s + 5×240s + 6×480s) and started immediately after the intravenous bolus injection of approximately 925 MBq of [^64^Cu]-ATSM. PET data were reconstructed using a LOR-RAMLA algorithm, with attenuation and scatter correction. Standard uptake value (SUV) images were calculated offline (with in-house written Matlab® quantification software) for each participant. PET and MRI data were normalized to Montreal Neurological Institute (MNI) space, the same geometric transformation was used, after the PET scan had been rigidly co-registered with the correspondent anatomical MRI scan (using SPM toolbox with default parameters; standard MRI brain was used when subject’s MRI were not available). An exploratory whole-brain and region-of-interest (ROI)-based analyses were performed. Data was exported for a set of distinct regions of interest including the cerebellum, basal ganglia and subregions (putamen, caudate and subthalamic nucleus) (regions known to be affected in HD patients). The SUV image was calculated per subject using the vessels uptake values as reference. Vessels uptake were defined as the average of the max uptake values over the first 3 minutes of the dynamic acquisition. Individual time-activity curves for the different ROIs and the uptake sum were determined during 10 to 26 minutes of acquisition.

### *In vivo* animal study

#### Animal’s characterization

YAC128 transgenic hemizygous (line HD53; 128 CAG repeats) and littermate non-transgenic wild-type (WT) mice (FVB/N background) with 3, 6, 9 and 12 months of age (mo) were used. All animals were generated from our local colony, with breeding couples gently provided by Dr. Michael Hayden (University of British Columbia, Vancouver, Canada). All animals were genotyped by common procedures as described in “Supplemental Methods” section. YAC128 mice exhibited an age-dependent gain of body weight (**Figure S1A**), as described previously^30,31^. All studies were carried out in accordance with the guidelines of the Institutional Animal Care and Use of Committee and the European Community directive (2010/63/EU) and protocols approved by the Faculty of Medicine, University of Coimbra (ORBEA_189_2018/11042018).

### [^64^Cu]-ATSM PET/MRI acquisitions

Animal PET acquisitions were performed by using the radiopharmaceutical [^64^Cu]-ATSM in YAC128 and WT mice at 3, 6, 9 and 12 mo. In all PET scans a prototype of a high-acceptance small-animal PET based on resistive plate chambers (RPC-PET) was used^32^. A mean activity of 418 ± 85 kBq/g was injected. The PET acquisition lasted for 60 minutes post injection. Volumetric images were reconstructed using OSEM algorithm and cubic voxel of 0.5 mm width. Two structural volumetric MRI T2 images were acquired per mouse using a MRI scan with the objective to facilitate the registration (geometric alignment) of the functional volumetric PET images with the volumetric MRI images, and thus allowing the segmentation of the regions of interest. A fiducial marker that can be viewed both in the PET and MRI imaging was placed in the mice bed. MRI imaging was performed after PET without moving the mice from the bed. Mice were kept anesthetized by isoflurane (1-2%) with 100% O_2_ with body temperature and respiration monitoring (SA Instruments SA, Stony Brook, USA). All MRI experiments were performed in a BioSpec 9.4T MRI scanner (Bruker Biospin, Ettlingen, Germany) with a volume head coil. High resolution morphological images were acquired with a 2D T2-weighted turbo RARE sequence and axial slice orientation. The sequence for fiducial marks had the following parameters: TR\/TE=8372\/33 ms, FOV=25.6×25.6 mm, acquisition matrix=256×256, averages=1, rare factor=8, echo spacing=11 ms, 80 axial continuous slices with 0.4 mm thick and acquisition time of 4m28s. The sequence for head mouse had the following parameters: TR\/TE=4500\/33 ms, FOV=25.6×25.6 mm, acquisition matrix=256×256, averages=8, rare factor=8, echo spacing=11 ms, 42 axial continuous slices with 0.4 mm thick and acquisition time of 19m12s.

### [^64^Cu]-ATSM uptake quantification

Based on the fiducial marker, the volumetric images obtained from the PET were manually registered by the volumetric MRI images using the 3D Slicer 4.4 software. The volumetric MRI images were manually segmented using the ITK-SNAP 2.2 software and the segmentation results applied to the registered volumetric PET images. The mean counts per mm^3^ in each ROI were computed and normalized by the activity injected per gram. Only the counts between 20- and 60-minutes post injection were considered.

### *In vitro* human and animal study

#### Cell culture

Fibroblasts from HD carriers and controls were generated from a small skin sample (about 3 mm diameter) from the abdominal region via a punch biopsy, as described previously^33^ (see “Supplemental Methods” for detailed description). Fibroblasts were cultured in DMEM medium (Gibco), supplemented with 15% FBS (Gibco) and 1% penicillin/streptomycin (Gibco) and maintained to a maximum of 15 passages.

### Isolation of functional mitochondria by Percoll gradient

Mice were weighted and then sacrificed by cervical dislocation and decapitation. Brains were removed from the skull and washed once in phosphate saline buffer (PBS) containing (in mM): 137 NaCl, 2.7 KCl, 1.8 KH_2_PO_4_, 10 Na_2_HPO_4_·2H_2_O, pH 7.4, followed by striatum and cortex dissection. Striatal and cortical mitochondrial-enriched fractions were then isolated using discontinuous Percoll density gradient centrifugation as previously described^34^. Briefly fresh striatal and cortical tissues were homogenized in ice-cold buffer (225 mM mannitol, 75 mM sucrose, 1 mM EGTA, 5 mM HEPES, pH 7.2/KOH, plus 1 mg/mL BSA) and centrifuged at 1,100 x*g* for 2 minutes at 4°C. The resulting supernatant was mixed with 80% Percoll (1 M sucrose, 50 mM HEPES, 10 mM EGTA, pH 7.0) and then carefully layered on the top of freshly made 10% Percoll (prepared from 80% Percoll) and centrifuged at 18,500 *xg* for 10 minutes at 4°C. Supernatant was discarded and the pellet was resuspended in 1 mL of washing buffer (250 mM sucrose, 5 mM HEPES-KOH, 0.1 mM EGTA, pH 7.2) and centrifuged again at 10,000 *xg* for 5 minutes at 4°C. Finally, the mitochondrial pellet was resuspended in ice-cold washing buffer and the amount of protein quantified by the Bio-Rad protein assay. Isolated mitochondria-enriched fractions were kept on ice for further functional analysis or frozen at −80°C.

### Bioenergetic assay

The fibroblasts oxidative phosphorylation and glycolytic profile were obtained by measuring the oxygen consumption rate (OCR) and extracellular acidification ratio (ECAR) on a Seahorse XF24 or XF96 apparatus. OCR was also analyzed in fresh striatal and cortical mitochondria isolated from YAC128 *versus* WT mice in coupling and uncoupling conditions by using a Seahorse XF24 apparatus (see “Supplemental Methods” for detailed description).

### Measurement of cellular and mitochondrial hydrogen peroxide levels

Mitochondrial or cellular hydrogen peroxide (H_2_O_2_) levels were measured in fibroblasts using mitochondria peroxy yellow 1 (MitoPY1) (Sigma) or Amplex Red reagent and horseradish peroxidase from the Amplex^®^ Red Catalase Assay Kit (Molecular Probes), respectively. H_2_O_2_ production by YAC128 and WT striatal and cortical mitochondria was measured by the Amplex® Red method. These procedures are detailed in “Supplemental Methods”.

### Transmission electron microscopy

For transmission electron microscopic (TEM) ultrastructural analyses fibroblasts were pelleted by centrifugation and striatum dissected out from 3 mo WT and YAC128 mice brain and processed as detailed in “Supplemental Methods”.

### Genomic DNA extraction and mitochondrial DNA copies

Genomic DNA was isolated from fibroblasts by means of high salt/ethanol precipitation. All DNA samples were considered pure when A260/A280 ratio was comprised between1.8–2.0. To quantify the average mitochondrial DNA (mtDNA) copy number of the fibroblasts the Absolute Human Mitochondrial DNA Copy Number Quantification qPCR Assay Kit (ScienCell Research Laboratories) was used according to manufacter protocol. Briefly, a mtDNA primer set recognizes and amplifies human mtDNA and another reference primer set recognizes and amplifies a 100 bp-long region on human chromosome 17 and serves as reference for data normalization. The reference genomic DNA sample with known mtDNA copy number serves as a reference for calculating the mtDNA copy number of target samples.

### Mitochondrial Ca^2+^ handling capacity

Mitochondrial Ca^2+^ uptake was measured fluorimetrically using the Ca^2+^-sensitive fluorescent dye Calcium Green 5N (150 nM; excitation and emission wavelengths of 506 nm and 532 nm, respectively) that exhibits an increase in fluorescence emission intensity upon binding to Ca^2+^ as described elsewhere^35^ with minor modifications^34^; thus, a decrease in the Calcium Green fluorescence indicates the capacity of mitochondria to handle Ca^2+^. Briefly, 5 μg of YAC128 and WT striatal or cortical isolated mitochondria was added to the incubation medium containing 125 mM KCl, 0.5 mM MgCl_2_, 3 mM KH_2_PO_4_, 10 mM Hepes, 10 μM EGTA, supplemented with 3 mM pyruvate, 1 mM malate, 3 mM succinate, 3 mM glutamate, 0.1 mM ADP and 1 μM oligomycin, pH 7.4. After a basal fluorescence record, six pulses of 10 μM CaCl_2_ were added every 4 minutes. Data are presented as traces of calcium handling capacity and plotted as extramitochondrial calcium (Extra mitoCa) for the third pulse of added calcium, as described in figure legend.

### Additional methods

See *Supplementary Materials* for more comprehensive description of the mentioned methods and details on genotyping, measurement of glutathione levels and glutathione peroxidase and glutathione reductase activities.

### Statistical analyses

Statistical computations were performed using GraphPad Prism version 7.0, GraphPad Software, La Jolla, CA, USA, and SPSS version 21.0 (IBM SPSS Statistics for Windows, IBM Corp). Statistical analysis was performed for individual subjects and among groups (controls, Pre-M and HD/manifest patients). For fibroblasts experiments at least three independent assays were performed for each experimental condition. Statistical significance was analysed using parametric test, two-way ANOVA, followed by Bonferroni post-hoc test and non-parametric test Kruskal Wallis followed by Dunn’s multiple comparison test. Correlations were done using the Spearman rank correlation coefficient (ρ). The power analysis approach to sample size determination was done using G power software on effect size (mean/standard deviation within groups) for 1–beta = 0.8 and alpha = 0.05. The results obtained from the animal’s experiments are expressed as the mean ± SEM of the number of replicates, considering the number of animals indicated in the figure legends. Comparisons among multiple groups were performed by two-way ANOVA followed by Tukey’s post-hoc test. Comparison between two populations, as described in figure legends was performed by nonparametric Mann Whitney U test. Significance was defined as p<0.05.

## RESULTS

### Clinical data

From the selected nine participants for this study, six were *HTT* gene mutation carriers, classified in premanifest and prodromal (Pre-M) and in HD manifest (**Table 1**). Pre-M HD carriers had no clinical motor signs or symptoms, based on standardized total motor score of Unified Huntington Disease Rating Scale (UHDRS)^36^. HD carriers had CAG repeats ranging from 42-52; the prodromal patient exhibited the longest repeat expansion and an anticipated phenotypic onset, compared to age-matched HD carriers. Manifest HD patients showed later symptom disease onset than Pre-M carriers. Furthermore, symptoms at onset varied from psychiatric (manifest HD patients) and cognitive (prodromal HD patient) symptoms.

### Increased accumulation of [^64^Cu]-ATSM in premanifest HD carriers

Whole brain and regional PET [^64^Cu]-ATSM retention in the SUV images was compared between groups using statistical parametric mapping and ROI analysis. Despite the reduced number of patients, the values obtained in the analysed areas of HD patients and controls were similar. Nonetheless, an apparent increased accumulation of [^64^Cu]-ATSM was observed in the Pre-M HD carriers, when compared to the control group. The exception was the caudate as the SUV decreased with disease severity, suggesting neuronal degeneration (**Figure 1A-C**). Spearman correlation analyses showed that [^64^Cu]-ATSM brain accumulation in whole brain and subthalamic nucleus (STN) directly correlates with CAG repeat number (ρ=0.709; p= 0.074; ρ=0.746, p=0.05, respectively) (**Figure 1D,E**).

**Figure 1:**
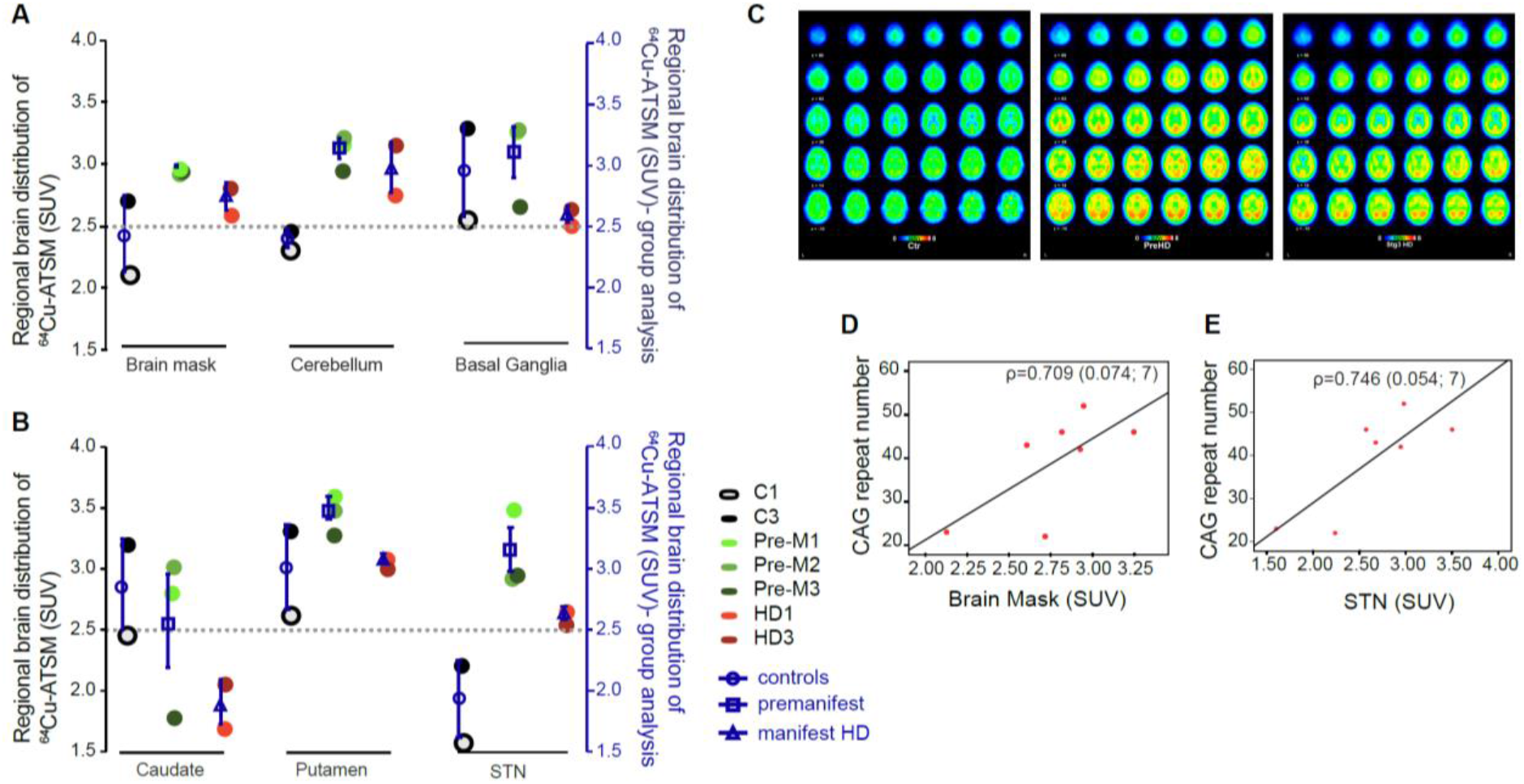
[^64^Cu]-ATSM brain accumulation in premanifest and manifest HD patients and correlation with CAG repeats. [^64^Cu]-ATSM brain PET imaging of HD patients and controls was evaluated in cerebellum and basal ganglia **(A)** and basal ganglia subregions: caudate, putamen and subthalamic nucleus (STN) **(B)** of controls **(C)**, premanifest (Pre-M) and manifest HD (HD) patient’s brains, as described in Methods section. (C) Representative PET imaging of the 3 groups analyzed: controls, premanifest and manifest HD patients; PET color scale indicates the uptake of the [^64^Cu]-ATSM normalized for cerebral blood vessels SUVs. Correlation between CAG repeat number and normalized SUV of [^64^Cu]-ATSM uptake from total brain **(D)** and uptake from putamen anterior **(E)**. Regional brain distribution of [^64^Cu]-ATSM **(A)(B)** was calculated as the sum of 3 time-points SUVs between 14 to 22 minutes acquisition time; mean ± SEM values of the groups are plotted on the secondary Y axis (blue). **(D)(E)** Correlation was performed using the Spearman correlation coefficient ρ (sig; n). SUV - standardized uptake values.

Enhanced [^64^Cu]-ATSM complex entrapment in an intracellular overreductive state observed in the majority of brain areas in Pre-M HD carriers can be caused by altered mitochondrial respiratory chain in patient’s brains.

### Altered mitochondrial function and redox deregulation in premanifest/prodromal HD carriers

Because [^64^Cu]-ATSM PET labelling has been attributed to overreduced intracellular state due to altered mitochondrial function and enhanced production of ROS ^24,25,27^, we assessed changes in mitochondrial function and ROS levels in human skin-derived fibroblasts obtained from controls and HD carriers (Pre-M and HD) detailed in **Table 1**.

Comparisons of oxygen consumption rates (OCR) between groups showed that basal and maximal respiration are enhanced in Pre-M *versus* controls accompanied by increased H^+^ leak (p<0.05) (**Figure 2A-D**). Contrarily, manifest HD patients demonstrated a general decrease in mitochondrial bioenergetic profile in basal (*vs* controls), maximal, spare respiration and H^+^ leak (*vs* Pre-M), p<0.05 (**Figure 2A-D**). No significant differences in ATP production (**Figure 2A-D**), ATP/ADP ratio or energy charge (**Figure S2**) were found in HD carriers, when compared to the control group. The observed increase in H^+^ leak in Pre-M HD carriers can be responsible for the increase in oxygen consumption without ATP synthesis.

**Figure 2:**
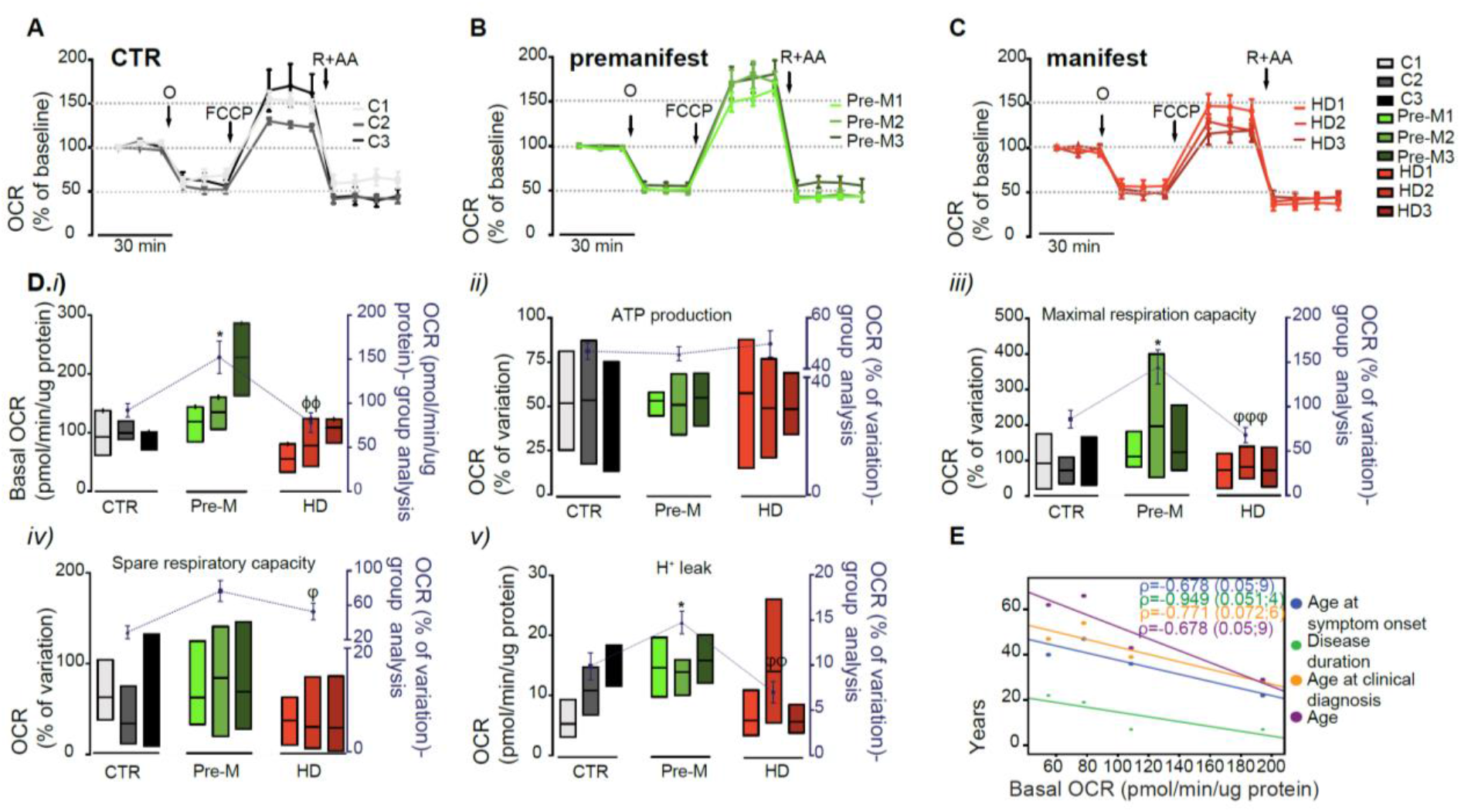
Oxygen consumption rate (OCR) in skin fibroblasts from premanifest and manifest HD carriers and controls. **(A-C)** OCR of cells treated with the different compounds. The mitochondrial inhibitors were sequentially injected into different ports of the Seahorse XF24 analyzer and the final concentrations of each were: 1 μM oligomycin (O), 0.3 μM FCCP (F), 1 μM rotenone and 1 μM antimycin A (R+A). **(D)** (i) Levels of basal OCR; (ii) Oxygen consumed for ATP generation through the complex V; (iii) Maximal respiration capacity; (iv) Spare respiratory capacity; (v) Component of OCR representing passive H^+^ leakage across the mitochondrial inner membrane were calculated as described in Methods section; individual data are presented as floating bars (min, max) with mean lines shown at least 3 independent experiments; group analysis represents the mean±SEM values (controls (CTR), premanifest (Pre-M) and manifest HD patients) and are plotted on the secondary Y axis (blue). **(E)** Correlation was performed using the Spearman correlation coefficient ρ (sig; n). Statistical analysis: One-Way ANOVA – post hoc Bonferroni’s multiple comparisons test. *p<0.05 (controls vs premanifest); ϕϕp<0.01 (controls *vs* HD manifest) and φp<0.05, φφp<0.01, φφφp<0.001 (premanifest *vs* HD manifest).

Interesting, we observed an inverse correlation between individual disease duration, age of diagnosis and onset and chronological age with basal OCR and a trend towards statistical significance. Longer disease duration (ρ=-0.949, p=0.051), more advanced age at symptom onset (ρ=-0.678, p=0.05) and at clinical diagnosis (ρ=-0.771, p=0.072) were accompanied by lower absolute values of basal respiration. Furthermore, older patients had significantly lower values of basal respiration (ρ=-0.678, p=0.05) (**Figure 2E**).

We also analysed glycolytic parameters by assessing extracellular acidification rates (ECAR) in human skin fibroblasts derived from the same individuals. Data showed no significant differences between groups, except for the glycolytic capacity of manifest HD patients that had lower absolute values, compared to controls (p<0.05) and Pre-M HD carriers (p<0.01) (**Figure S3**).

The levels of ROS are largely determined by mitochondrial dysfunction, triggering an overreduction state that may lead to oxidative stress involved in [^64^Cu]-ATSM accumulation. Thus, the relative levels of mitochondrial and cellular hydrogen peroxide (H_2_O_2_) were measured in human skin fibroblasts using MitoPY1 and Amplex Red, respectively (**Figure 3**). We found significantly higher levels of cellular H_2_O_2_ in fibroblasts derived from Pre-M HD carriers, which remained constant along disease progression (**Figure 3A**). Interestingly, a significant rise in basal levels of mitochondrial H_2_O_2_ (mito-H_2_O_2_) were observed in more advanced stages of HD only (**Figure 3B**). Following complex III inhibition with myxothiazol (3 μM), the levels of mito-H_2_O_2_ increased along disease severity, being significantly higher in cells derived from manifest HD patients *versus* controls (p<0.05) (**Figure 3C**). Correlation analysis showed augmented cellular H_2_O_2_ levels in HD carriers with longer CAG repeat number (ρ=0.769, p=0.05), and a lower uptake of [^64^Cu]-ATSM in the caudate was associated with increased cellular H_2_O_2_ levels (ρ= −0.714, p=0.071) (**Figure 3D,E**).

**Figure 3:**
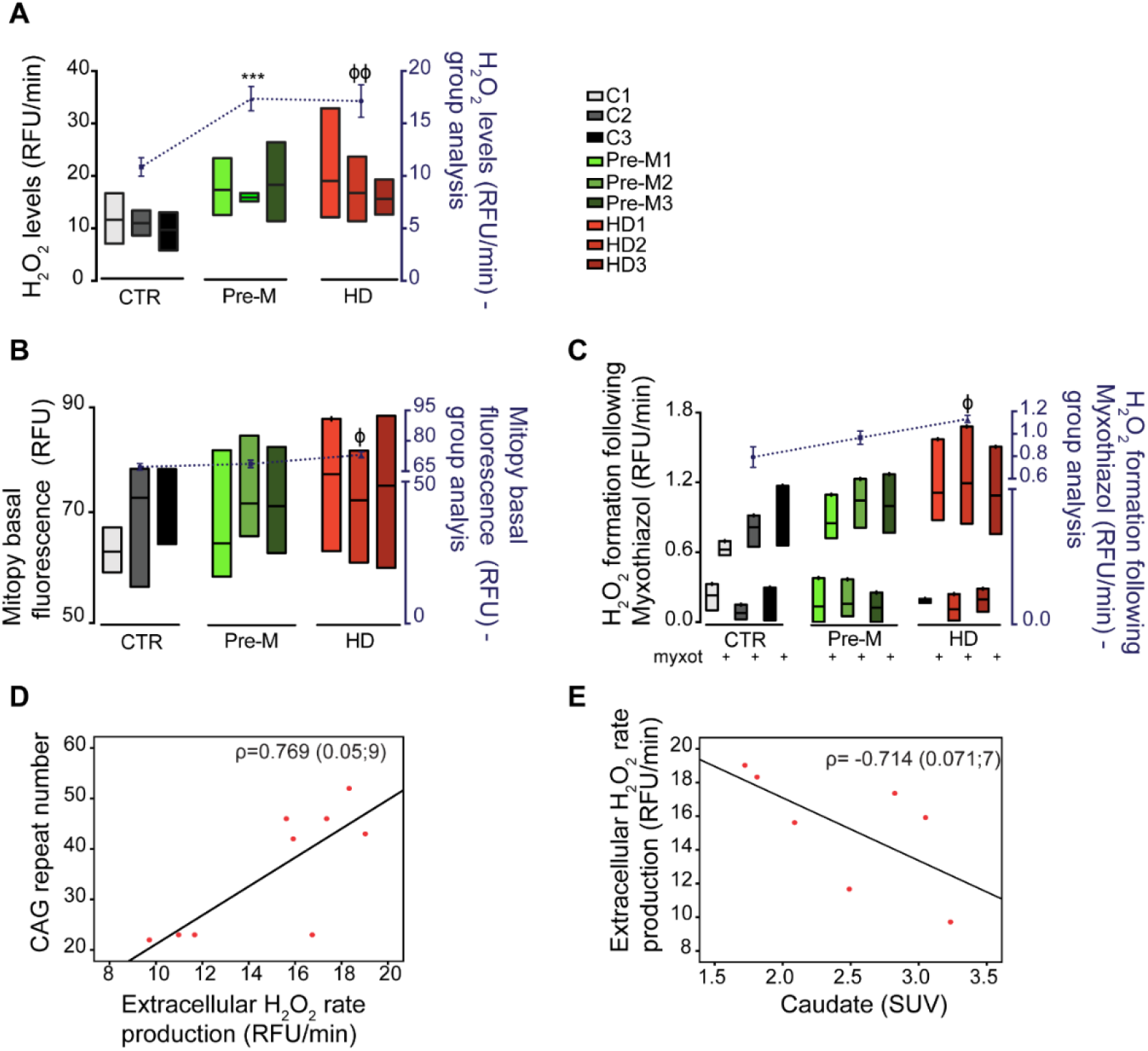
Cellular and mitochondrial ROS levels in skin fibroblasts from HD patient’s and controls - correlation with CAG repeats and [^64^Cu]ATSM uptake in caudate. Cellular **(A)** and basal mitochondrial **(B)** H_2_O_2_ production were measured as described in Methods section and was plotted as bars graph. **(C)** Time-dependent H2O2 production in the mitochondria after stimulation with complex III inhibitor-myxothiazol (3 μM). Individual data are presented as floating bars (min, max) with mean lines shown at least 3 independent experiments; group analysis represents the mean±SEM values (controls (CTR), pre-symptomatic (Pre-M) and manifest HD patients) and are plotted on the secondary Y axis (blue). **(D)(E)** Correlation was performed using the Spearman correlation coefficient ρ (sig; n). Statistical analysis: non-parametric Kruskal-Wallis test followed by Dunn’s Multiple Comparison Test. ***p<0.001 (controls *vs* premanifest); and ϕp<0.05, ϕϕp<0.01 (controls *vs* HDs).

These data indicate early increased mitochondrial respiration linked to H^+^ leak and redox deregulation that is heightened in manifest HD.

### Premanifest and prodromal HD carriers show decreased roundness and increased mitochondrial DNA copies correlated to higher [^64^Cu]-ATSM uptake

Because mitochondrial dysfunction has been linked to changes in mitochondrial morphology, we analysed the morphometric parameters of mitochondria that were individually traced from TEM images obtained in human skin fibroblasts (**Figure 4A**). We measured the major mitochondrial shape factors, namely area, circularity, Feret’s diameter, perimeter, form factor, and aspect ratio (**Figure 4B**). Mitochondria from Pre-M HD carriers show decreased circularity, are more elongated (higher Feret’s diameter), have more complex and branched networks and larger aspect ratio (length/width ratio) when compared with the control or manifest HD patients (**Figure 4A,B; Suppl. Table 1**) (p<0.05). However, increased circularity can be observed in manifest HD patient’s fibroblasts, when compared to the controls (p<0.05), indicating that mitochondria from fibroblasts of Pre-M HD carriers are less fragmented.

**Figure 4:**
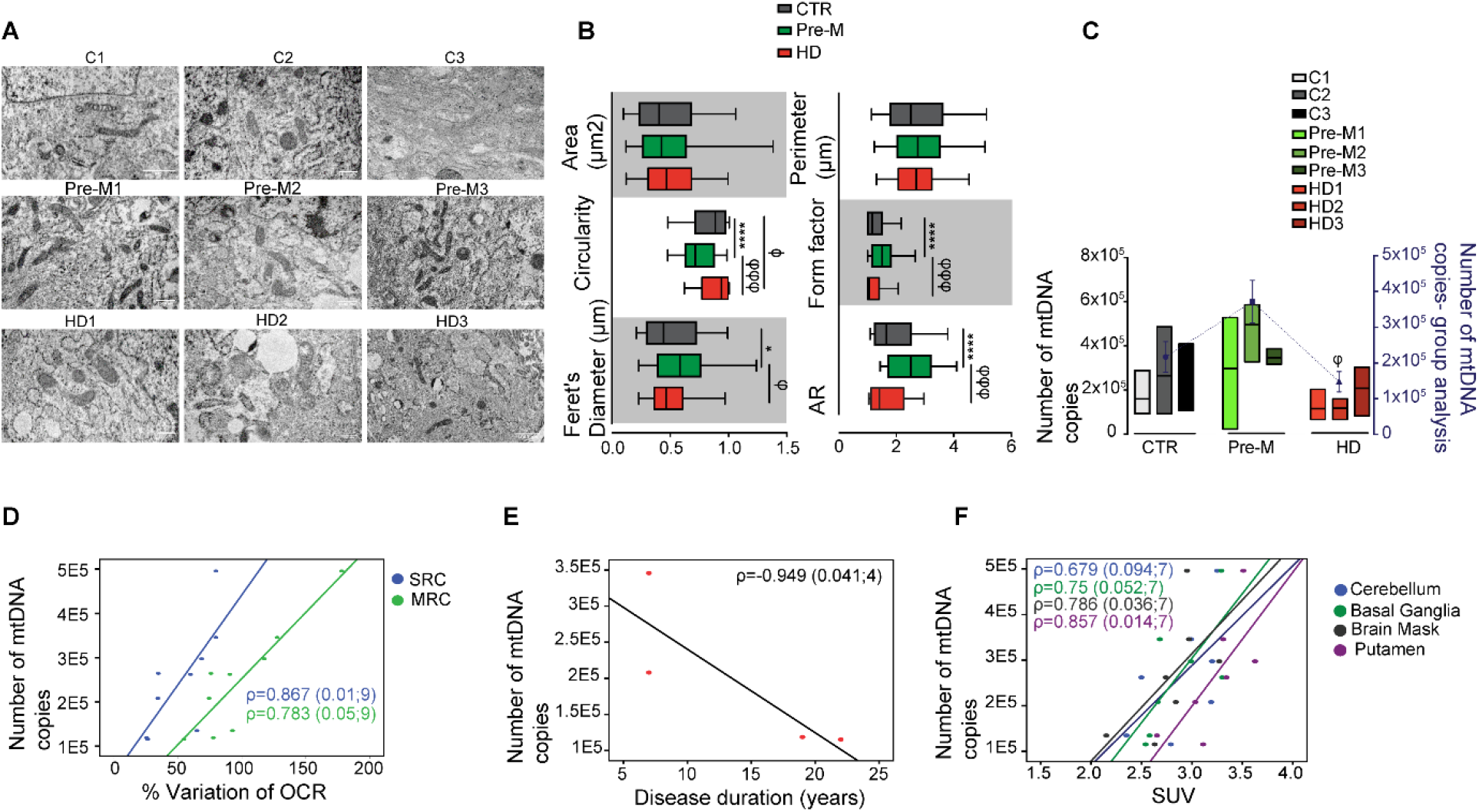
Mitochondrial morphological features and DNA copy number in fibroblasts derived from premanifest and manifest HD carriers and controls - correlation with [^64^Cu]-ATSM uptake, disease duration and OCR. **(A)** Transmission electron microscopy (TEM) representative images of mitochondrial ultrastructure. **(B)** Analysis of morphological parameters for mitochondria area, circularity, Feret’s diameter, perimeter, form factor and aspect ratio made using ImageJ software. The results are expressed as the mean±SEM of mitochondria counted by images (n=9-26) from a total of n=29-122 mitochondria analysed. Bars represent the median and interquartile range [percentiles 5-95] for each parameter. Statistical analysis: ANOVA followed post-hoc Bonferroni test: ****p<0.0001 (controls *vs* pre-symptomatic); ϕp<0.05, (controls *vs* HDs) and φp<0.05, φφφp<0.001 (pre-M *vs* HDs). **(C)** Number of mitochondrial DNA copies quantified as described in Methods section. Individual data are presented as floating bars (min, max) with mean lines shown at least 3 independent experiments; group analysis represents the mean±SEM values (controls (CTR), premanifest (Pre-M) and manifest HD carriers) and are plotted on the secondary Y axis (blue). Statistical analysis: non-parametric Kruskal-Wallis test followed by Dunn’s Multiple Comparison Test. φp<0.05 (pre-M *vs* HDs). **(D-F)** Correlation was performed using the Spearman correlation coefficient ρ (sig; n). Scale bar: 2000 nm.

Fibroblasts from Pre-M HD carriers show a trend for increased number of mtDNA copies, whilst manifest HD patients have reduced mtDNA copy number when compared to cells from Pre-M carriers (p<0.05) (**Figure 4C**). Furthermore, we observed that human skin fibroblasts with higher number of mitochondrial DNA copies have significantly greater spare (ρ=0.867, p=0.01) and maximal respiratory capacity (ρ=0.783, p=0.05). In accordance, patients with longer disease duration present a lower number of mtDNA copies (ρ=-0.949, p=0.041). Interestingly, enhanced mtDNA copy number was observed in patients with augmented [^64^Cu]-ATSM uptake in whole brain (ρ=0.786, p=0.036), cerebellum (ρ=0.679, p=0.094), basal ganglia (ρ=0.75, p=0.052) and putamen (ρ=0.857, p=0.014). These data suggest that Pre-M HD carriers showing higher number of mtDNA copies in peripheral cells also have higher accumulation of [^64^Cu]-ATSM in selected brain areas, which may reflect a mitochondrial overreduced state.

### Enhanced [^64^Cu]-ATSM brain accumulation in YAC128 mouse brain

In order to verify whether changes in [^64^Cu]-ATSM brain accumulation correlate with mitochondrial changes in HD brain affected areas, we further analysed *in vivo* [^64^Cu]-ATSM accumulation by PET and studied mitochondrial fractions isolated from the striatum and cortex of YAC128 mice along aging.

*In vivo* age-dependent analysis of intracellular overreductive status was evaluated in YAC128 *versus* wild-type mice over the progression of the disease, at 3 (pre-symptomatic), 6 (early-symptomatic) and 9 and 12 (symptomatic) months of age (mo), through the analysis of [^64^Cu]-ATSM accumulation in the striatum (**Figure 5A,B**) and frontal cortex (**Figure 5A,C**). Statistical analysis by two-way ANOVA revealed a significant genotype and age-dependent [^64^Cu]-ATSM accumulation in striatum (F1,35)=12.07; p=0.0014) and (F3,35)=36.67; p<0.0001), respectively, evidencing enhanced total accumulation of the radioligand in YAC128 compared to WT striatum (**Figure 5B*i***). *In vivo* data obtained from the cortex also demonstrated an age-dependent effect F(3,38)=12.61; p<0.0001) and a slight, but significant, genotype effect in [^64^Cu]-ATSM accumulation F(1,18)=4.192; p=0.0476 (**Figure 5C*i***). Accordingly, when all ages were merged for both YAC128 and WT striatum and cortex, a significant increase in overall radioligand accumulation was observed, indicating increased overreductive status in these two YAC128 mouse brain areas. In addition, when [^64^Cu]-ATSM accumulation in YAC128 mouse striatum or cortex were replotted in relation to WT mice, a significant increase in the radioligand accumulation was observed in the striatum of 6 mo (p<0.05) and a trend at 12 mo (p=0.0571) (**Figure 5B*ii***), but not in the cortex (**Figure 5C*ii***).

**Figure 5:**
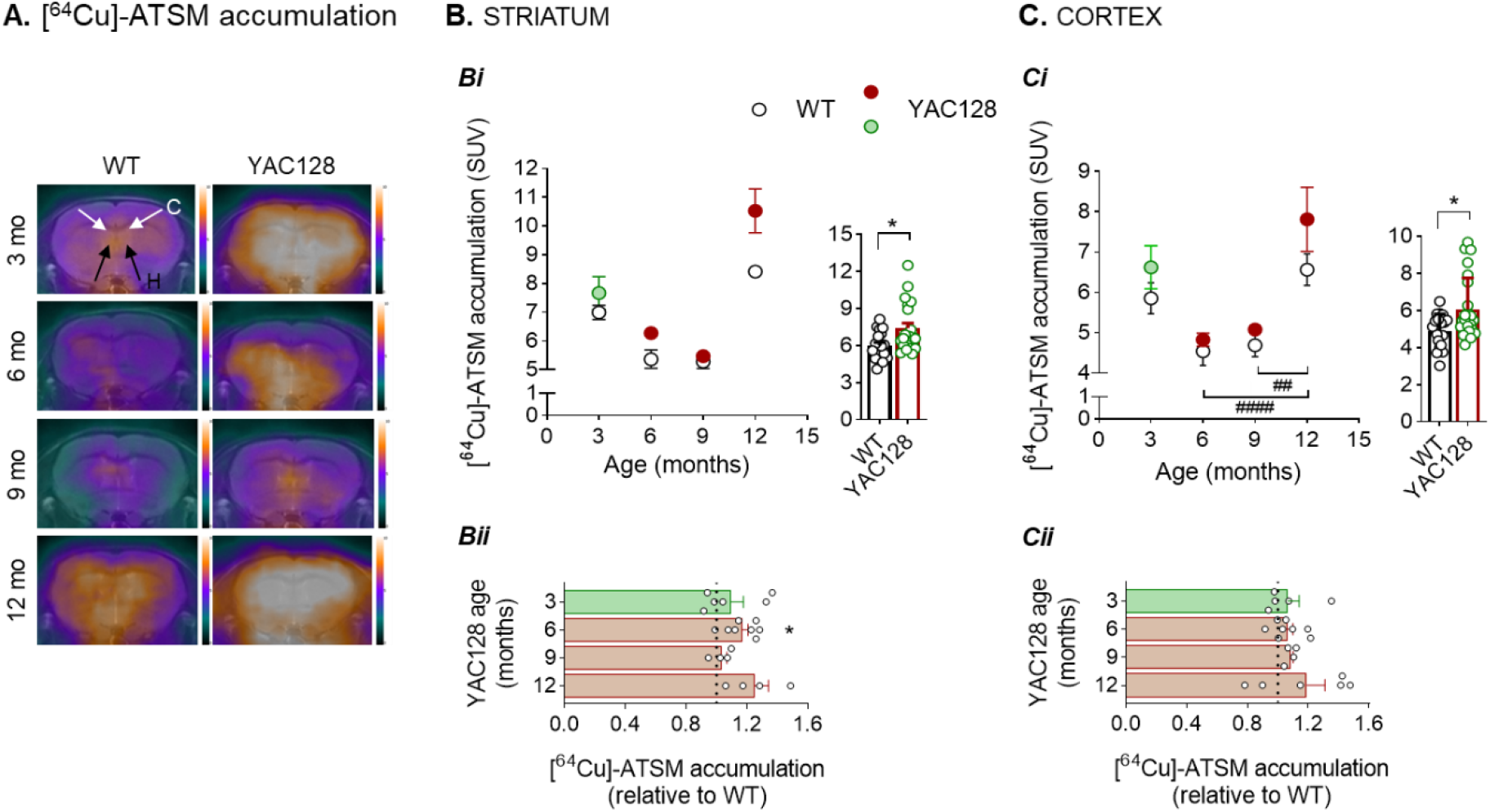
Age-dependent [^64^Cu]-ATSM brain accumulation in YAC128 and WT mouse striatum and cortex. [^64^Cu]-ATSM PET and MRI overlapped axial images of brain YAC128 and WT mice where PET color scale indicates the uptake (number of counts per mm^3^) of the [^64^Cu]-ATSM normalized by the injected activity, mass of the mouse and sensibility of the PET equipment. White arrows point to frontal cortex and black arrows to striatum (**A**). [^64^Cu]-ATSM accumulation was plotted for striatum (**B**) and cortex (**C**) at 3, 6, 9 and 12 mo YAC128 vs WT mice, as described in Methods section and replotted as total radioligand accumulation for all ages of each genotype (insets **B*i* and *Ci*** for striatum and cortex, respectively) and in percentage of WT for each age for striatum (***Bii***) and cortex (**C*ii***). Statistical analysis: *p<0.05 when compared with 6 mo WT mitochondria by nonparametric Mann-Whitney U test. SUV - standardized uptake values.

### Increased oxygen consumption rate, complexes II and III activities and oxidative status in striatal mitochondria derived from pre-symptomatic YAC128 mice

The respiratory activity in striatal and cortical isolated mitochondrial-enriched fractions derived from YAC128 and WT mouse brain striatum and cortex was analysed by determining OCR and complexes activities (**Figures 6 and S4**). Notably, both wild-type HTT and mHTT were shown to associate with cortical- and striatal-enriched isolated mitochondria derived from WT and YAC128 mice, respectively (**Figure S1C**).

**Figure 6:**
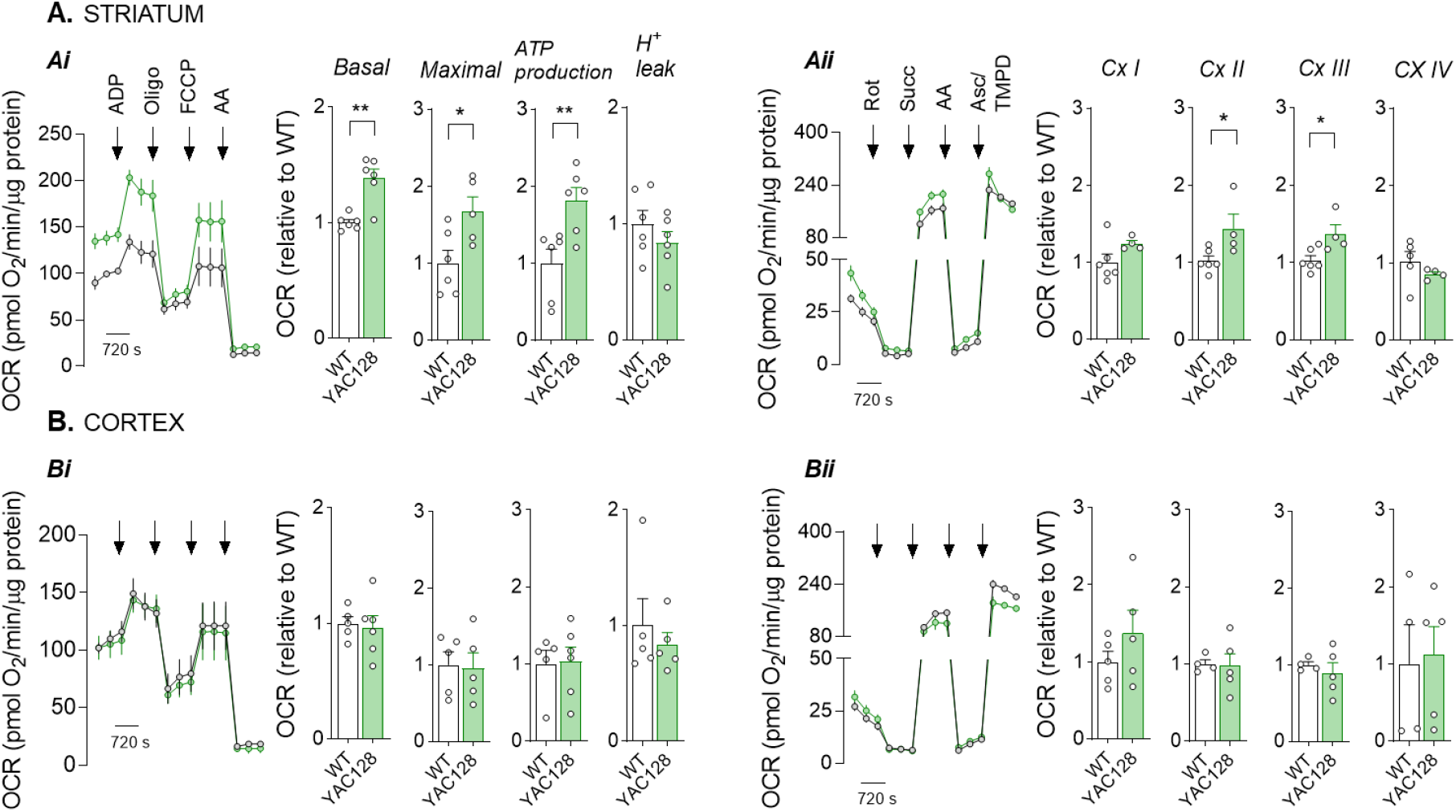
Oxygen consumption rates (OCR) in 3 mo YAC128 mouse striatal and cortical mitochondria. Mitochondrial respiration was measured using the Seahorse flux analyzer in 3 mo YAC128 and WT striatal (**A**) and cortical (**B**) isolated mitochondria. Levels of respiratory coupling (**A*i* and B*i***; for striatal and cortical mitochondria, respectively) were analysed in MAS containing 10 mM succinate plus 2 μM rotenone under sequentially injection of mitochondrial inhibitors and substrates (final concentration: 4 mM ADP; 2.5 μg/mL oligomycin; 4 μM FCCP and 4 μM antimycin A) as shown in representative traces and basal respiration and maximal respiration, ATP production and H^+^ calculated as described in Methods section. The activity of mitochondrial respiratory chain complexes (**A*ii* and B*ii***; for striatal and cortical mitochondria, respectively) was analyzed in uncoupling conditions performed in MAS containing 4 μM FCCP, 10 mM pyruvate and 2 mM malate. Mitochondrial inhibitors and substrates were sequentially injected (final concentration: 2 μM rotenone, 10 mM succinate, 4 μM antimycin A and 10 mM ascorbate/100 μM TMPD) as shown in the representative traces and Cx I-IV activities calculated as described in Methods section. Data are the mean ± SEM of experiments performed in independent mitochondrial preparations obtained from 3-7 mice from each genotype, run in duplicates or triplicates. Statistical analysis: *p<0.05, **p<0.01 when compared with WT mitochondria, by nonparametric Mann-Whitney U test.

Coupling experiments performed under Cx II feeding (in the presence of succinate) and Cx I inhibition (with rotenone) in striatal mitochondria obtained from 3 mo (pre-symptomatic) YAC128 mice showed increased basal and maximal respiration and ATP production, but unchanged H^+^ leak, (**Figure 6A*i***), when compared to WT mice. In order to further explain these changes in OCR, we analyzed the electron flow through mitochondrial respiratory chain, to evaluate Cx I-IV activities under conditions of direct mitochondrial feeding (pyruvate, to generate acetyl-CoA) plus malate (to activate the TCA cycle) in FCCP-induced uncoupled state. In agreement with results obtained in **Figure 6A*i***, striatal mitochondria obtained from 3 mo YAC128 mice exhibited a significant increase in Cx II and III activities, but unaltered activities in Cx I and IV (**Figure 6A*ii***), when compared with WT mouse mitochondria. No significant changes in mitochondrial-coupled respiration parameters or Cx I-IV activities were observed in cortical mitochondria (at 3, 6, 9 and 12 mo) (**Figure 6 B*i* and B*ii*; Figure S4B*i* and S4B*ii***) or striatal mitochondria obtained from older YAC128 mice (at 6, 9 and 12 mo) (**Figure S4A*i* and S4A*ii***). Data suggest a striatum-specific increase in mitochondrial respiratory chain activity and Cx II and III activity in mitochondria from pre-symptomatic YAC128 mice.

As in human fibroblasts, we further analyzed the levels of mitochondrial H2O2 produced by *ex vivo* striatal- and cortical mitochondrial-enriched fractions isolated from pre-symptomatic (3 mo) and symptomatic (6, 9 and 12 mo) YAC128 mice and age-matched WT mice (**Figure 7A*i* and 7A*ii***). Results evidenced a small but significant increase in H2O2 production by striatal mitochondria from both pre-(3 mo; p<0.05) and late-symptomatic (12 mo; p<0.01) mice (**Figure 7A*i***) and by cortical mitochondria from late symptomatic YAC128 mice, at 12 mo (p<0.05) (**Figure 7A*ii***) without significant changes in intermediate ages (6 and 9 mo) for both brain areas.

**Figure 7:**
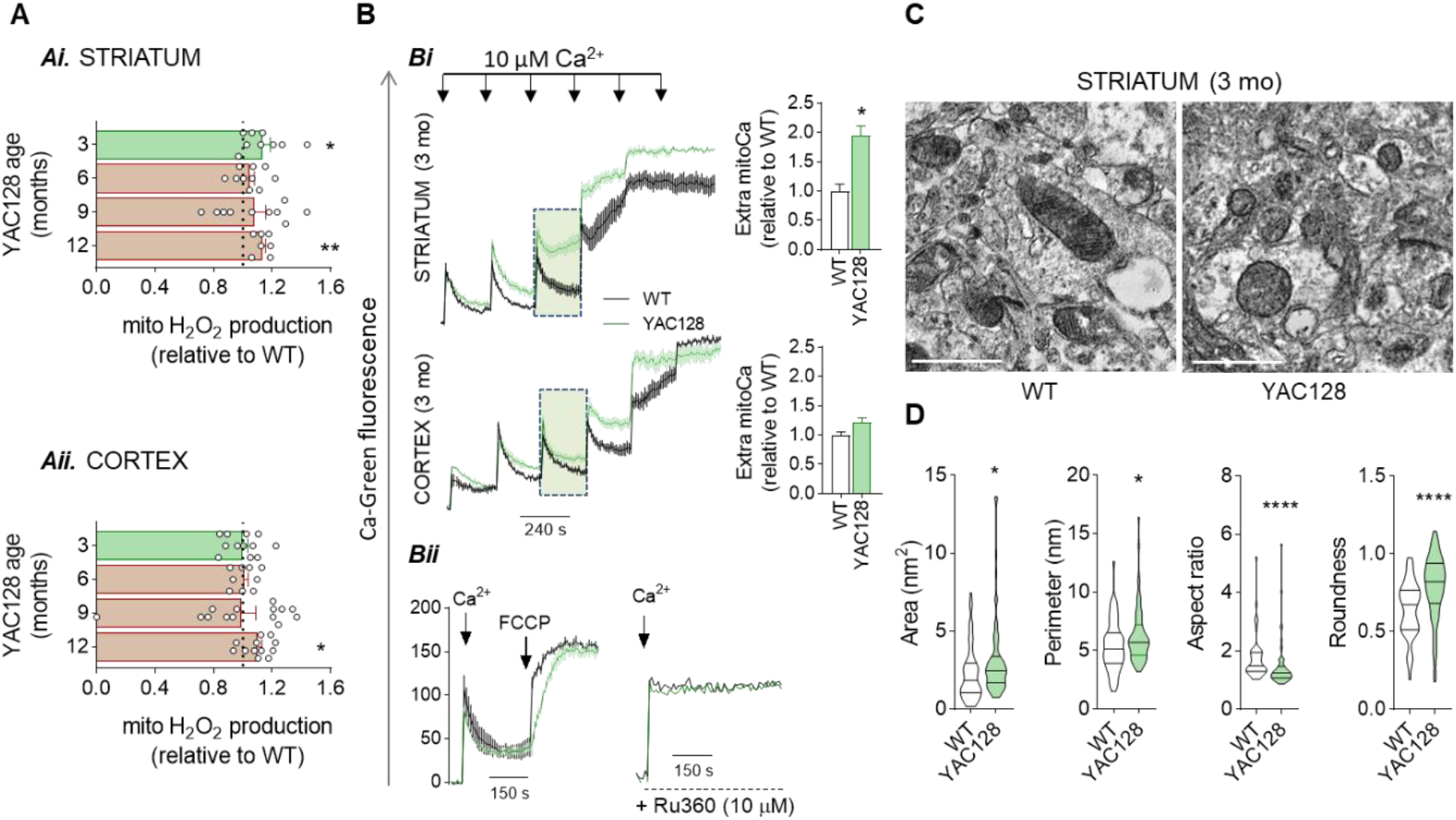
Mitochondrial ROS levels, Ca^2+^ handling, and mitochondrial ultrastructure in YAC128 mouse striatal and cortical mitochondria. H_2_O_2_ production was measured in functional isolated mitochondria (**A**) derived from 3, 6, 9 and 12 mo YAC128 and WT mice striatum (**A*i***) and cortex (**A*ii***) using the Amplex Red assay as described in Methods section. The capacity to handle Ca^2+^ was evaluated in striatal and cortical mitochondria obtained from 3 mo YAC128 and WT mice (**B**) by using Calcium Green-5N fluorescent probe. Pulses of 10 μM Ca^2+^ were applied to raise intramitochondrial Ca^2+^ concentrations until the Ca^2+^ retention capacity was reached as shown by the upward deflection of the traces (**B*****i**)*. The Ca^2+^ handling capacity for striatal and cortical mitochondria at the third Ca^2+^ pulse is detailed in the dashed green window and presented as extra-mitochondrial Ca^2+^ (area under the curve) (**B*i, inset***). The effect of the mitochondrial uncoupler FCCP and the pharmacological inhibition of the mitochondrial calcium uniporter (MCU; Ru360) are also depicted (**B*ii***). Electron micrographs were obtained from 3 mo YAC128 and WT mice striatum and mitochondrial morphology presented for area, perimeter, aspect ratio and roundness as evaluated by Fiji-imageJ software (**C**). Over 62 individual mitochondria were assessed in independent preparations from 6 independent TEM images from WT and YAC128 mice (scale bar: 1000 nm). Data are the mean ± SEM of experiments performed in independent mitochondrial preparations obtained from 3-16 mice from each genotype run in triplicates to quadruplicates. Statistical analysis: *p<0.05, **p<0.01, ****p<0.0001 when compared with WT mitochondria, by nonparametric Mann-Whitney U test.

**Graphical Abstract/Figure 8:**
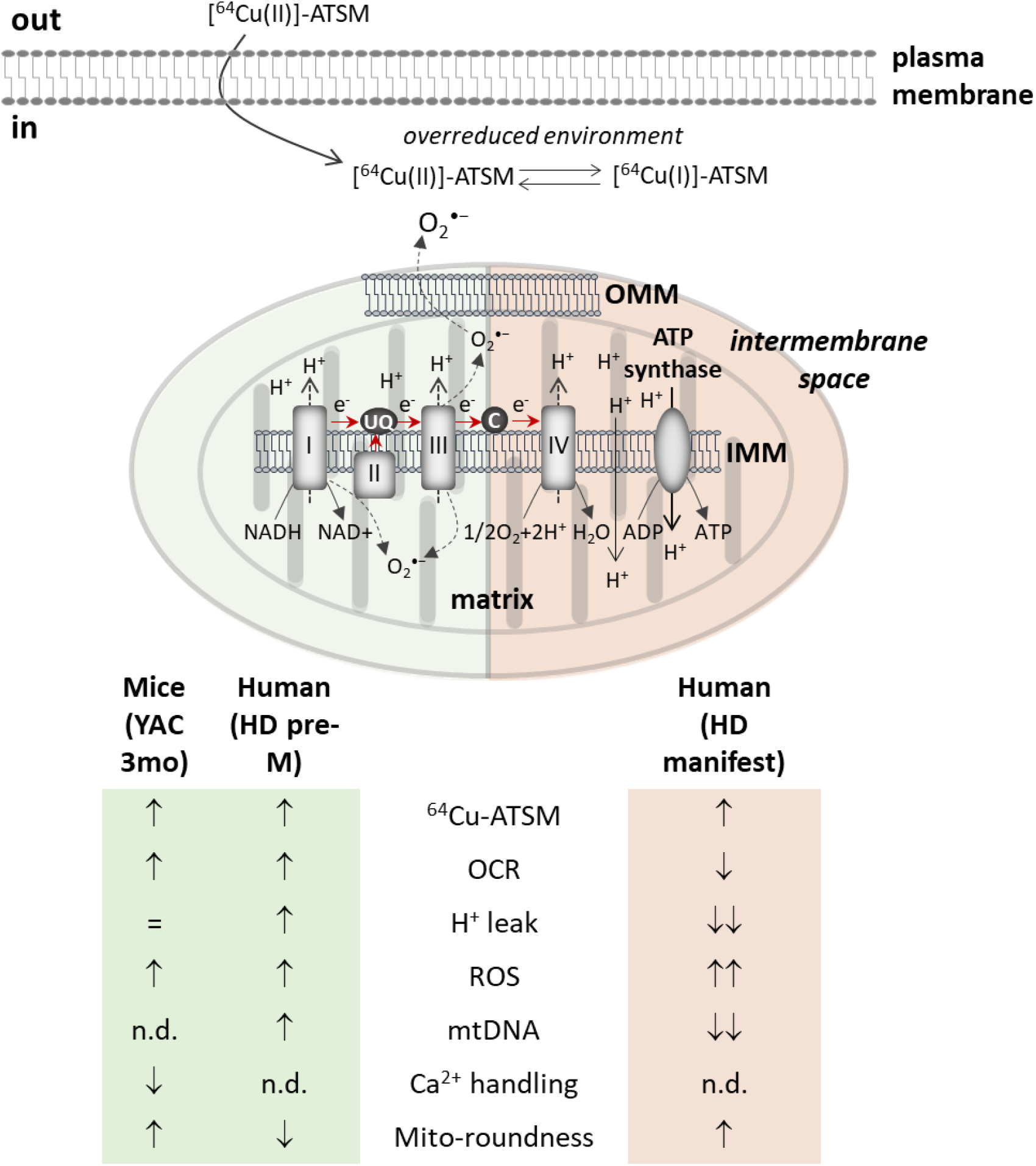
Overview of brain [^64^Cu]-ATSM retention and mitochondrial abnormalities in cells from Huntington’s disease (HD) carriers, at premanifest and manifest disease stages, and in presymptomatic YAC128 mice. In premanifest/prodromal (pre-M) HD carriers, HD manifest and YAC128 mice at pre-symptomatic (3 months of age, mo) stages an electron rich environment induced by increased ROS production and impaired electron transport chain can result in an overreduced state. Under these conditions, Cu(II)-ATSM is reduced to Cu(I)-ATSM after entering the cell, turning the complex unstable, and free copper-64 is trapped and accumulated in the intracellular space. Interestingly, in both peripheral cells from pre-M HD carriers and striatal mitochondria from pre-symptomatic (3 mo) YAC128 mice a similar increase in OCR is observed; in YAC128 mouse mitochondria this occurs along with enhanced activity of mitochondrial complexes, decreased Ca^2+^ handling and enhanced mitochondrial roundness. In contrast, cells from HD manifest patients show higher mitochondrial dysregulation, coincident with reduced mitochondrial DNA (mtDNA) copy number.

Elevated mito-H_2_O_2_ levels were not accompanied by compromised antioxidant defenses in YAC128 mice at 3 and 9 mo, since no significant changes were observed in mitochondrial acetyl(K68)SOD2/SOD2 or SOD2 protein levels in both striatum (**Figure S5A*i***) and cortex (**Figure S5B*i***), reduced (GSH) and oxidized (GSSG) glutathione total levels in striatum (**Figure S5Aii and S5A*iii***, respectively) or cortex (**Figure S5B*ii* and S5B*iii***, respectively), nor glutathione peroxidase and reductase activities in both striatum (**Figure S5A*iv* and S5A*v***, respectively) and cortex (**Figure S5B*iv* and S5B*v***, respectively), despite significant decreased glutathione reductase protein levels in mitochondria from 3 mo YAC128 mouse striatum (p<0.05) (**Figure S5A*vi***), but not in cortex (**Figure S5B*vi***). Similarly, data evidenced no differences in antioxidants at 12 mo in striatum or cortex (data not shown).

### Early Ca^2+^ deregulation and altered ultrastructure in YAC128 mouse striatal mitochondria

Because altered mitochondrial Ca^2+^ retention may serve as an indicator of mitochondrial impairment, we next evaluated Ca^2+^ handling in striatal and cortical mitochondria-enriched fractions isolated from 3 mo YAC128 and WT mice. Mitochondrial Ca^2+^ uptake was monitored after sequential additions of 10 μM Ca^2+^ to raise intramitochondrial Ca^2+^ concentration in 3 mo striatal (**Figure 7B*i*, upper panel**) and cortical (**Figure 7B*i*, lower panel**) mitochondria. Under these conditions, Ca^2+^ was sequentially taken up by both striatal and cortical mitochondria (**Figure 7B*i***), however the response to the 3rd pulse of Ca^2+^ produced a decrease in Ca^2+^ retention in striatal (p<0.05; **Figure 7B*i*, upper panel**, *detailed in dashed green window*), but not in cortical (**Figure 7B*i*, lower panel**, *detailed in dashed green window*) YAC128 mitochondria, when compared to WT mitochondria, as evaluated by extramitochondrial Ca^2+^ (Extra-mitoCa) quantification; the following pulse of Ca^2+^ in striatal (**Figure 7B*i*, upper panel**), but not in cortical (**Figure 7B*i*, lower panel**) mitochondria exceeded the mitochondrial Ca^2+^ retention capacity, with a subsequent abrogation of Ca^2+^ uptake as represented by upward deflection of the trace, indicating the opening of the mitochondrial permeability transition pore (mPTP). Our data highlight defects in Ca^2+^ handling in striatal mitochondria derived from pre-symptomatic YAC128 mice, exhibiting lower Ca^2+^ thresholds for mPTP opening after repeated Ca^2+^ loads. Ca^2+^ taken up by WT and YAC128 striatal-derived mitochondria was shown to be released through a FCCP-induced “releasable pool” (**Figure 7B*ii***) and prevented by pre-incubation with the mitochondrial Ca^2+^ uniporter (MCU) inhibitor, Ru360 (**Figure 7B*ii***), indicating that Ca^2+^ retention occurred through the MCU in polarized mitochondria. Similar results were observed in cortical mitochondria (data not shown).

In order to investigate YAC128 mouse striatal mitochondrial morphology, electron micrographs obtained by TEM of 3 mo YAC128 and WT mice striatum were assessed using quantitative measures for mitochondrial morphology (**Figure 7C**). Over 154 individual mitochondria were assessed for area, perimeter, aspect ratio and roundness (**Figure 7D**). YAC128 mouse striatal mitochondria presented increased area and perimeter along with decreased aspect ratio and a more roundness appearance, whereas WT mitochondria appeared ellipsoidal. These observations are consistent with the loss of normal mitochondrial appearance *in situ*, suggesting increased mitochondrial fragmentation despite higher area, when compared to WT mitochondria (**Figure 7D**), supporting dysfunctional YAC128 striatal mitochondria.

## DISCUSSION

This work is the first to demonstrate evidence of higher accumulation of [^64^Cu]-ATSM in premanifest and prodromal (Pre-M) HD carriers and in an early-stage HD animal model in different brain regions. The exception was the caudate, a region reported to be affected in the prodromal phase and showing progressive atrophy along the course of the disease due to neuronal loss^37–39^. High levels of [^64^Cu]-ATSM retention appear to be indicative of an overreduction state^27^. In accordance, our results show that fibroblasts from Pre-M HD carriers and striatal mitochondria derived from pre-symptomatic YAC128 mice exhibit increased mitochondrial hydrogen peroxide levels, which is unrelated with decreased antioxidant levels or activity, but rather with an apparently compensatory enhanced activity of mitochondrial respiratory chain and altered morphology.

Classically, disease burden is defined based on CAG repeat length and age. In this study we show a correlation between imaging of the whole brain and particularly the STN and the extent of CAG repeats. The STN is a key component of the basal ganglia and several reports, in animal models, argue that STN dysfunction and neuronal loss precede cortico-striatal abnormalities in HD, which might contribute to motor impairment, and explain the greater accumulation of [^64^Cu]-ATSM in Pre-M individuals^40,41^.

The excessively overreduced microenvironment found in Pre-M HD carriers was coupled to an overall increase in mitochondrial bioenergetics, including higher values of basal OCR and maximal respiration in skin fibroblasts. Nevertheless, adenine nucleotide levels remained unaffected, probably due an increase in basal OCR uncoupled from ATP synthesis, measured as H^+^ leak. In skin fibroblasts, we also found lower values of basal OCR in HD patients correlating with earlier age of clinical diagnosis and age of onset, with longer disease duration and increased age. Additionally, a lower mitochondrial network fragmentation occurred concomitantly with a tendency for upregulation in mtDNA copy number in peripheral cells from pre-M HD carriers. Importantly, mtDNA levels are increased in younger pre-M carriers with shorter disease duration and correlate with enhanced mitochondrial bioenergetics and increased accumulation of [^64^Cu]-ATSM, probably as a reflex of the redox state already documented at this stage.

In YAC128 mice, increased [^64^Cu]-ATSM retention was mostly evident in striatum of 6 and 12 mo and less manifested in the cortex. Interestingly, striatal isolated mitochondria (but not cortical mitochondria) derived from pre-symptomatic (3 mo) YAC128 mice exhibited not only increased levels of H_2_O_2_hydrogen peroxide, but also enhanced activity of complexes II and III, linked to increased basal and maximal OCR and ATP production. Increased complexes activity and H_2_O_2_ levels were recently observed by us in isolated YAC128 mouse striatal mitochondria at 3 mo^12^. The increase in mitochondrial ATP generation in striatal mitochondria contrasts with human fibroblasts showing unchanged ATP levels due to H^+^ leakage. In addition, striatal mitochondria from pre-symptomatic YAC128 mice showed defects in Ca^2+^ handling and altered mitochondrial morphology, suggesting deregulated mitochondrial function despite the increase in mitochondrial respiratory activity, which precedes the increase in [^64^Cu]-ATSM retention, at 6 mo. Altered striatal mitochondrial function observed in this study implicates enhanced susceptibility of this brain region, consistently with previous reports showing a highly susceptibility of the striatum to impaired mitochondrial oxidative phosphorylation^42^ and age-dependent striatal volumetric changes defined by us through longitudinal structural imaging in the YAC128 mouse model^43^. Unchanged OCR in YAC128 mouse mitochondria at 6-12 mo may be accounted for by alternative metabolic profile and/or decreased striatal energy demand, as demonstrated by enhanced levels of creatine and phosphocreatine, determined by spectroscopy (1H-MRS)^43^.

Several studies have established a link between [^64^Cu]-ATSM accumulation, mitochondrial dysfunction, impairment of mitochondrial respiratory chain, due e.g. hypoxia, and related oxidative stress^26,27,44^, suggesting that dysfunctional mitochondrial electron transport chain causes an overreduced state due to ROS generation. Furthermore, in a patient with MELAS (mitochondrial encephalopathy, lactic acidosis, and stroke-like episode), caused by a mitochondrial mutation, [^62^Cu]-ATSM accumulation reflected changes in regional oxidative stress^26^. Increased ROS production and mitochondrial dysfunction were also described in Parkinson’s disease and amyotrophic lateral sclerosis patients, as evaluated by PET using [^62^Cu]-ATSM and further associated with disease severity in both diseases^24,25,45^.

In HD, mHTT is widely expressed in all body cells turning energy metabolism studies in peripheral cells important in the search for disease biomarkers. A previous study stressed that fibroblasts from healthy donor and HD patients exhibited different profiles if stratified according to motor function, psychiatric and functional capacity. Fibroblasts obtained from manifest HD patients displayed compromised cell growth and reduced ATP production, accompanied by decreased mitochondrial activity, which was indicative of respiratory chain defects^15^. Of interest, the age at onset seems to be a crucial element alongside the CAG expansion to define the bioenergetic profile of fibroblasts of HD patients^14^. Concordantly, our data suggest that HD patients suffering for longer periods of time have highly compromised mitochondrial function. In these cells, a slight increase in mitochondrial ROS was reported, despite a significant elevation in antioxidant enzymes (SOD2 and GR)^15^ while other authors described a decrease in catalase activity in HD fibroblasts^22^. Impaired antioxidant defenses, namely a decrease in GPx and SOD1 activities, were also observed in erythrocytes from HD patients^46^. Previous studies performed in our lab showed increased mitochondrial-driven ROS in HD human cybrids^18^ and mouse striatal cells expressing full-length mHTT^20,21,47,48^. An increase in ROS was also found in the brain striatum and cortex of symptomatic R6/1 ^49^, R6/2 HD mouse models^50,51^ and in YAC128 embryonic fibroblasts and fibroblasts derived from HD patients^52^. Despite enhanced mitochondrial production of H2O2, our data showed unchanged antioxidant activity in cortical and striatal mitochondria at all ages, implicating that decreased antioxidant profile does contribute for altered redox changes in HD mouse brain.

Mitochondrial function has a high impact on organelle morphology, defined by a regulated balance between fission and fusion events, since loss of bioenergetic capacity results in an inability to maintain the mitochondrial network organization^53^. In our study, skin fibroblast mitochondria from Pre-M HD carriers became larger, whereas circularity decreased, when compared to healthy controls and manifest patients. These ultrastructural characteristics suggest a promotion of mitochondrial fusion events that might be required to maximize oxidative phosphorylation. Conversely, striatal mitochondria from pre-symptomatic YAC128 mice, retaining mHTT interaction, exhibited increased roundness, as revealed by ultrastructure analysis. Such opposing results might be determined by cell type–specific vulnerability of human peripheral cells or isolated organelle derived from mouse brain cells (neurons and glia) in the most affected brain area in HD. Accordingly, fragmented mitochondria was previously reported in many HD cell models, including peripheral cells (e.g.^54^). Mitochondrial fragmentation was also observed in rat cortical neurons expressing mHTT and in fibroblasts from manifest HD patients and in YAC128 HD mouse model, before the appearance of neurological deficits and mHTT aggregates^55^; in striatum of YAC128 mice, an increase in the number of small mitochondria and decreased cristae surface area were also detected^55^.

Decreased ability to take up Ca^2+^ through the MCU was further observed in striatal mitochondrial-enriched fractions from pre-symptomatic YAC128 mice, when compared to cortical mitochondria, again underlying a highly dysfunctional organelle derived from the striatum. Defects on mitochondrial Ca^2+^ handling were previously demonstrated in lymphoblast mitochondria from HD patients and brain mitochondria from YAC128 mice^56^, isolated mitochondria from transgenic HD rats with 51 CAG repeats^57^ and in mutant mouse striatal cell line^58,59^. Concordantly, in striatal mitochondria obtained from rat brain, lower calcium levels consistently evoked a permeability transition easily than in cortical mitochondria ^60^. Conversely, an increase in Ca^2+^ loading capacity in forebrain (brain minus the cerebellum)-derived mitochondria from R6/2 and YAC128 mice, but not in organelle isolated from Hdh150 knock-in mice, was observed^16^. More recently, Pellman and co-authors evidenced a large Ca^2+^ uptake capacity in synaptic and nonsynaptic mitochondria isolated from 2 and 12 mo YAC128 mice brain, when compared with YAC18 or FVB/NJ mice^35^. Indeed, changes in mitochondrial Ca^2+^ handling have not been consistent in HD models, particularly when studying isolated brain mitochondria, which might be related with different methodologies of isolating the organelle and/or assessing ion changes, as well as the isolated brain tissue.

The number of mtDNA copies has a crucial role in cellular metabolism and bioenergetics. The mechanisms responsible for regulation of mtDNA copy number in HD pathology are still under discussion. Our study has revealed that Pre-M HD carriers harbor a higher number of mtDNA copies; interestingly, a strong and significant correlation was observed with enhanced mitochondrial bioenergetics and [^64^Cu]-ATSM uptake. Contradictory conclusions may be dependent on the cell type, as shown by a significantly higher mtDNA/nDNA copy number in leukocytes, but lower in fibroblasts from manifest HD patients^61^. Moreover, mtDNA copy number was increased before disease onset in leukocytes of HD mutation carriers and declined after disease onset^62^.

Of relevance, this study provides novel and important data suggesting a compensatory mechanism in premanifest and prodromal HD carriers and pre-symptomatic HD mice model to overcome the early insults, particularly the underlying oxidative stress. Compensatory mechanisms at early stages of age-related brain diseases, as determined in human and mouse, have been described elsewhere^63,64^. Reports of upregulation in the expression of mitochondrial-encoded genes in blood samples of mild cognitively impaired patients, compared to AD patients and healthy controls, suggested a compensatory-like mechanism at the early stages of disease^65,66^.

Overall, this study provides novel data regarding major redox changes and mitochondrial deregulation at early symptomatic stages in HD, as revealed by *in vivo* PET analysis showing accumulation of [^64^Cu]-ATSM; this is a powerful tool to improve our insights into oxidative stress linked to mitochondrial dysfunction, sustaining an early oxidant status. The major limitation of this study includes the small number of participants. Further studies with a larger number of participants and at different disease timepoints are necessary to validate human data. Nonetheless, the overlapping results obtained in mitochondrial function and oxidative stress in pre-symptomatic stages in YAC128 mice supports the changes reported in patients. Moreover, data in mouse striatum reinforces an increased susceptibility of this brain region in HD. Thus, potential benefits of early therapeutic interventions based on mitochondrial-targeted compounds aiming to regularize mitochondrial function and reduce ROS generation, particularly in the striatum, are anticipated to help to slowdown HD progression.

## ACKNOWLEDGMENTS

This work was funded by Mantero Belard Neuroscience prize 2013 (1^st^ Edition), supported by Santa Casa da Misericórdia de Lisboa (SCML), Portugal, ‘FLAD Life Science 2020’ prize, funded by ‘Fundação Luso-Americana para o Desenvolvimento’ (FLAD), Portugal, FEDER through “Programa Operacional Factores de Competitividade – COMPETE” and “Fundação para a Ciência e a Tecnologia” (FCT), project references: UID/NEU/04539/2013; PEst-C/SAU/LA0001/2013-2014 and by the European Regional Development Fund (ERDF), through the Centro 2020 Regional Operational Programme: project CENTRO-01-0145-FEDER-000012-HealthyAging2020, the COMPETE 2020 - Operational Programme for Competitiveness and Internationalisation, and the Portuguese national funds via FCT – Fundação para a Ciência e a Tecnologia, projects UIDB/04539/2020 and UIDP/04539/2020.

Ferreira IL was supported by Mantero Belard Neuroscience prize 2013 (1st Edition), SCML post-doctoral fellowship and FCT postdoctoral fellowship SFRH/BPD/108493/2015; Mota SI and Laço M were supported by FCT postdoctoral fellowships SFRH/BPD/99219/2013 and SFRH/BPD/91811/2012, respectively. We acknowledge Dr. Mónica Zuzarte, for fibroblasts and mouse brain TEM analyses at the ‘Laboratório de Bio-imagem de Alta Resolução’ of the Faculty of Medicine of the University of Coimbra.

## List of abbreviations

ADP: Adenosine diphosphate
ATP: Adenosine triphosphate
BSA: Bovine serum albumin
CAG: Cytosine-Adenine-Guanine
EGTA: Ethylene glycol tetra-acetic acid
ETC: Electron transport chain
FCCP: Carbonyl cyanide p-trifluoromethoxyphenylhydrazone
GPx: Glutathione peroxidase
GRed: Glutathione reductase
GSH: Glutathione, reduced form
GSSG: Glutathione, oxidized form
HD: Huntington’s disease
H_2_O_2_: Hydrogen peroxide
HTT: Huntingtin
MCU: Mitochondrial calcium uniporter
mHTT: Mutant huntingtin
MIM: Mitochondrial inner membrane
mmp: Mitochondrial transmembrane potential
mPTP: Mitochondrial permeability transition pore
NaF: Sodium fluoride
NEM: N-ethylmaleimide
OCR: Oxygen consumption rate
PET: Positron emission tomography
ROS: Reactive oxygen species
SDS: Sodium dodecyl sulphate
SDS-PAGE: SDS polyacrylamide gel electrophoresis
SOD: Superoxide dismutase
STN: Subthalamic nucleus
YAC: Yeast artificial chromosome

## REFERENCES

1. Group THsDCR. A novel gene containing a trinucleotide repeat that is expanded and unstable on Huntington’s disease chromosomes. The Huntington’s Disease Collaborative Research Group. Cell. Mar 26 1993;72(6):971–83. doi:10.1016/0092-8674(93)90585-e

2. Damiano M, Galvan L, Deglon N, Brouillet E. Mitochondria in Huntington’s disease. Biochimica et biophysica acta. Jan 2010;1802(1):52–61. doi:10.1016/j.bbadis.2009.07.012

3. Costa V, Scorrano L. Shaping the role of mitochondria in the pathogenesis of Huntington’s disease. The EMBO journal. Apr 18 2012;31(8):1853–64. doi:10.1038/emboj.2012.65

4. Naia L, Ferreira IL, Ferreiro E, Rego AC. Mitochondrial Ca(2+) handling in Huntington’s and Alzheimer’s diseases - Role of ER-mitochondria crosstalk. Biochemical and biophysical research communications. Feb 19 2017;483(4):1069–1077. doi:10.1016/j.bbrc.2016.07.122

5. Brennan WA, Bird ED, Aprille JR. Regional mitochondrial respiratory activity in Huntington’s disease brain. J Neurochem. Jun 1985;44(6):1948–50. doi:10.1111/j.1471-4159.1985.tb07192.x

6. Browne SE, Bowling AC, MacGarvey U, et al. Oxidative damage and metabolic dysfunction in Huntington’s disease: selective vulnerability of the basal ganglia. Ann Neurol. May 1997;41(5):646–53. doi:10.1002/ana.410410514

7. Gu M, Gash MT, Mann VM, Javoy-Agid F, Cooper JM, Schapira AH. Mitochondrial defect in Huntington’s disease caudate nucleus. Ann Neurol. Mar 1996;39(3):385–9. doi:10.1002/ana.410390317

8. Damiano M, Diguet E, Malgorn C, et al. A role of mitochondrial complex II defects in genetic models of Huntington’s disease expressing N-terminal fragments of mutant huntingtin. Human molecular genetics. Oct 1 2013;22(19):3869–82. doi:10.1093/hmg/ddt242

9. Hamilton J, Pellman JJ, Brustovetsky T, Harris RA, Brustovetsky N. Oxidative metabolism in YAC128 mouse model of Huntington’s disease. Human molecular genetics. Sep 1 2015;24(17):4862–78. doi:10.1093/hmg/ddv209

10. Hamilton J, Brustovetsky T, Brustovetsky N. Oxidative metabolism and Ca(2+) handling in striatal mitochondria from YAC128 mice, a model of Huntington’s disease. Neurochemistry international. Oct 2017;109:24–33. doi:10.1016/j.neuint.2017.01.001

11. Hamilton J, Pellman JJ, Brustovetsky T, Harris RA, Brustovetsky N. Oxidative metabolism and Ca2+ handling in isolated brain mitochondria and striatal neurons from R6/2 mice, a model of Huntington’s disease. Human molecular genetics. Jul 1 2016;25(13):2762–2775. doi:10.1093/hmg/ddw133

12. Naia L, Ly P, Mota SI, et al. The Sigma-1 Receptor Mediates Pridopidine Rescue of Mitochondrial Function in Huntington Disease Models. Neurotherapeutics. Apr 2021;doi:10.1007/s13311-021-01022-9

13. Silva AC, Almeida S, Laço M, et al. Mitochondrial respiratory chain complex activity and bioenergetic alterations in human platelets derived from pre-symptomatic and symptomatic Huntington’s disease carriers. Mitochondrion. Nov 2013;13(6):801–9. doi:10.1016/j.mito.2013.05.006

14. Gardiner SL, Milanese C, Boogaard MW, et al. Bioenergetics in fibroblasts of patients with Huntington disease are associated with age at onset. Neurol Genet. Oct 2018;4(5):e275. doi:10.1212/NXG.0000000000000275

15. Jędrak P, Mozolewski P, Węgrzyn G, Więckowski MR. Mitochondrial alterations accompanied by oxidative stress conditions in skin fibroblasts of Huntington’s disease patients. Metab Brain Dis. 12 2018;33(6):2005–2017. doi:10.1007/s11011-018-0308-1

16. Oliveira JM, Jekabsons MB, Chen S, et al. Mitochondrial dysfunction in Huntington’s disease: the bioenergetics of isolated and in situ mitochondria from transgenic mice. Journal of neurochemistry. Apr 2007;101(1):241–9. doi:10.1111/j.1471-4159.2006.04361.x

17. Oliveira JM, Chen S, Almeida S, et al. Mitochondrial-dependent Ca2+ handling in Huntington’s disease striatal cells: effect of histone deacetylase inhibitors. J Neurosci. Oct 2006;26(43):11174–86. doi:10.1523/JNEUROSCI.3004-06.2006

18. Ferreira IL, Nascimento MV, Ribeiro M, et al. Mitochondrial-dependent apoptosis in Huntington’s disease human cybrids. Experimental neurology. Apr 2010;222(2):243–55. doi:10.1016/j.expneurol.2010.01.002

19. Ferreira IL, Cunha-Oliveira T, Nascimento MV, et al. Bioenergetic dysfunction in Huntington’s disease human cybrids. Experimental neurology. Sep 2011;231(1):127–34. doi:10.1016/j.expneurol.2011.05.024

20. Ribeiro M, Rosenstock TR, Cunha-Oliveira T, Ferreira IL, Oliveira CR, Rego AC. Glutathione redox cycle dysregulation in Huntington’s disease knock-in striatal cells. Free radical biology & medicine. Nov 15 2012;53(10):1857–67. doi:10.1016/j.freeradbiomed.2012.09.004

21. Ribeiro M, Rosenstock TR, Oliveira AM, Oliveira CR, Rego AC. Insulin and IGF-1 improve mitochondrial function in a PI-3K/Akt-dependent manner and reduce mitochondrial generation of reactive oxygen species in Huntington’s disease knock-in striatal cells. Free radical biology & medicine. Sep 2014;74:129–44. doi:10.1016/j.freeradbiomed.2014.06.023

22. del Hoyo P, García-Redondo A, de Bustos F, et al. Oxidative stress in skin fibroblasts cultures of patients with Huntington’s disease. Neurochem Res. Sep 2006;31(9):1103–9. doi:10.1007/s11064-006-9110-2

23. Sawa A, Wiegand GW, Cooper J, et al. Increased apoptosis of Huntington disease lymphoblasts associated with repeat length-dependent mitochondrial depolarization. Nat Med. Oct 1999;5(10):1194–8. doi:10.1038/13518

24. Ikawa M, Okazawa H, Kudo T, Kuriyama M, Fujibayashi Y, Yoneda M. Evaluation of striatal oxidative stress in patients with Parkinson’s disease using [62Cu]ATSM PET. Nucl Med Biol. Oct 2011;38(7):945–51. doi:10.1016/j.nucmedbio.2011.02.016

25. Ikawa M, Okazawa H, Tsujikawa T, et al. Increased oxidative stress is related to disease severity in the ALS motor cortex: A PET study. Neurology. May 2015;84(20):2033–9. doi:10.1212/WNL.0000000000001588

26. Ikawa M, Okazawa H, Arakawa K, et al. PET imaging of redox and energy states in stroke-like episodes of MELAS. Mitochondrion. Apr 2009;9(2):144–8. doi:10.1016/j.mito.2009.01.011

27. Yoshii Y, Yoneda M, Ikawa M, et al. Radiolabeled Cu-ATSM as a novel indicator of overreduced intracellular state due to mitochondrial dysfunction: studies with mitochondrial DNA-less ρ0 cells and cybrids carrying MELAS mitochondrial DNA mutation. Nucl Med Biol. Feb 2012;39(2):177–85. doi:10.1016/j.nucmedbio.2011.08.008

28. Reilmann R, Leavitt BR, Ross CA. Diagnostic criteria for Huntington’s disease based on natural history. Mov Disord. Sep 15 2014;29(11):1335–41. doi:10.1002/mds.26011

29. Matarrese M, Bedeschi P, Scardaoni R, et al. Automated production of copper radioisotopes and preparation of high specific activity [(64)Cu]Cu-ATSM for PET studies. Applied radiation and isotopes: including data, instrumentation and methods for use in agriculture, industry and medicine. Jan 2010;68(1):5–13. doi:10.1016/j.apradiso.2009.08.010

30. Pouladi MA, Xie Y, Skotte NH, et al. Full-length huntingtin levels modulate body weight by influencing insulin-like growth factor 1 expression. Human molecular genetics. Apr 15 2010;19(8):1528–38. doi:10.1093/hmg/ddq026

31. Lopes C, Ribeiro M, Duarte AI, et al. IGF-1 intranasal administration rescues Huntington’s disease phenotypes in YAC128 mice. Molecular neurobiology. Jun 2014;49(3):1126–42. doi:10.1007/s12035-013-8585-5

32. Martins P, Blanco A, Crespo P, et al. Towards very high resolution RPC-PET for small animals. Journal of Instrumentation. 2014;9(C10012)doi:10.1088/1748-0221/9/10/C10012

33. Onofre I, Mendonça N, Lopes S, et al. Fibroblasts of Machado Joseph Disease patients reveal autophagy impairment. Sci Rep. 06 2016;6:28220. doi:10.1038/srep28220

34. Ferreira IL, Carmo C, Naia L, I Mota S, Cristina Rego A. Assessing Mitochondrial Function in In Vitro and Ex Vivo Models of Huntington’s Disease. Methods Mol Biol. 2018;1780:415–442. doi:10.1007/978-1-4939-7825-0_19

35. Pellman JJ, Hamilton J, Brustovetsky T, Brustovetsky N. Ca(2+) handling in isolated brain mitochondria and cultured neurons derived from the YAC128 mouse model of Huntington’s disease. J Neurochem. Aug 2015;134(4):652–67. doi:10.1111/jnc.13165

36. Group. THS. Unified Huntington’s Disease Rating Scale: reliability and consistency. Mov Disord. Mar 1996;11(2):136–42. doi:10.1002/mds.870110204

37. Hobbs NZ, Barnes J, Frost C, et al. Onset and progression of pathologic atrophy in Huntington disease: a longitudinal MR imaging study. AJNR Am J Neuroradiol. Jun 2010;31(6):1036–41. doi:10.3174/ajnr.A2018

38. Harrington DL, Long JD, Durgerian S, et al. Cross-sectional and longitudinal multimodal structural imaging in prodromal Huntington’s disease. Mov Disord. 11 2016;31(11):1664–1675. doi:10.1002/mds.26803

39. Novak MJ, Seunarine KK, Gibbard CR, et al. White matter integrity in premanifest and early Huntington’s disease is related to caudate loss and disease progression. Cortex. Mar 2014;52:98–112. doi:10.1016/j.cortex.2013.11.009

40. Atherton JF, McIver EL, Mullen MR, Wokosin DL, Surmeier DJ, Bevan MD. Early dysfunction and progressive degeneration of the subthalamic nucleus in mouse models of Huntington’s disease. Elife. 12 2016;5doi:10.7554/eLife.21616

41. Callahan JW, Abercrombie ED. Relationship between subthalamic nucleus neuronal activity and electrocorticogram is altered in the R6/2 mouse model of Huntington’s disease. J Physiol. Aug 2015;593(16):3727–38. doi:10.1113/JP270268

42. Pickrell AM, Fukui H, Wang X, Pinto M, Moraes CT. The striatum is highly susceptible to mitochondrial oxidative phosphorylation dysfunctions. The Journal of neuroscience: the official journal of the Society for Neuroscience. Jul 6 2011;31(27):9895–904. doi:10.1523/JNEUROSCI.6223-10.2011

43. Petrella LI, Castelhano JM, Ribeiro M, et al. A whole brain longitudinal study in the YAC128 mouse model of Huntington’s disease shows distinct trajectories of neurochemical, structural connectivity and volumetric changes. Human molecular genetics. Jun 15 2018;27(12):2125–2137. doi:10.1093/hmg/ddy119

44. Donnelly PS, Liddell JR, Lim S, et al. An impaired mitochondrial electron transport chain increases retention of the hypoxia imaging agent diacetylbis(4-methylthiosemicarbazonato)copperII. Proc Natl Acad Sci U S A. Jan 2012;109(1):47–52. doi:10.1073/pnas.1116227108

45. Neishi H, Ikawa M, Okazawa H, et al. Precise Evaluation of Striatal Oxidative Stress Corrected for Severity of Dopaminergic Neuronal Degeneration in Patients with Parkinson’s Disease: A Study with 62Cu-ATSM PET and 123I-FP-CIT SPECT. Eur Neurol. 2017;78(3-4):161–168. doi:10.1159/000479627

46. Chen CM, Wu YR, Cheng ML, et al. Increased oxidative damage and mitochondrial abnormalities in the peripheral blood of Huntington’s disease patients. Biochemical and biophysical research communications. Jul 27 2007;359(2):335–40. doi:10.1016/j.bbrc.2007.05.093

47. Ribeiro M, Silva AC, Rodrigues J, Naia L, Rego AC. Oxidizing effects of exogenous stressors in Huntington’s disease knock-in striatal cells--protective effect of cystamine and creatine. Toxicological sciences: an official journal of the Society of Toxicology. Dec 2013;136(2):487–99. doi:10.1093/toxsci/kft199

48. Oliveira AM, Cardoso SM, Ribeiro M, Seixas RS, Silva AM, Rego AC. Protective effects of 3-alkyl luteolin derivatives are mediated by Nrf2 transcriptional activity and decreased oxidative stress in Huntington’s disease mouse striatal cells. Neurochemistry international. Dec 2015;91:1–12. doi:10.1016/j.neuint.2015.10.004

49. Perez-Severiano F, Santamaria A, Pedraza-Chaverri J, Medina-Campos ON, Rios C, Segovia J. Increased formation of reactive oxygen species, but no changes in glutathione peroxidase activity, in striata of mice transgenic for the Huntington’s disease mutation. Neurochemical research. Apr 2004;29(4):729–33. doi:10.1023/b:nere.0000018843.83770.4b

50. Sadagurski M, Cheng Z, Rozzo A, et al. IRS2 increases mitochondrial dysfunction and oxidative stress in a mouse model of Huntington disease. The Journal of clinical investigation. Oct 2011;121(10):4070–81. doi:10.1172/JCI46305

51. Tabrizi SJ, Workman J, Hart PE, et al. Mitochondrial dysfunction and free radical damage in the Huntington R6/2 transgenic mouse. Annals of neurology. Jan 2000;47(1):80–6. doi:10.1002/1531-8249(200001)47:1<80::aid-ana13>3.3.co;2-b

52. Wang JQ, Chen Q, Wang X, et al. Dysregulation of mitochondrial calcium signaling and superoxide flashes cause mitochondrial genomic DNA damage in Huntington disease. The Journal of biological chemistry. Feb 1 2013;288(5):3070–84. doi:10.1074/jbc.M112.407726

53. Lopes C, Tang, Y., Anjo, S.I., Manadas, B., Onofre, I., De Almeida, L.P., Daley, G.Q., Schlaeger, T.M., and Rego, A.C. Mitochondrial and Redox Modifications in Huntington Disease Induced Pluripotent Stem Cells Rescued by CRISPR/Cas9 CAGs Targeting. Frontiers in Cell and Developmental Biology. 2020;8doi:10.3389/fcell.2020.576592

54. Mihm MJ, Amann DM, Schanbacher BL, Altschuld RA, Bauer JA, Hoyt KR. Cardiac dysfunction in the R6/2 mouse model of Huntington’s disease. Neurobiology of disease. Feb 2007;25(2):297–308. doi:10.1016/j.nbd.2006.09.016

55. Song W, Chen J, Petrilli A, et al. Mutant huntingtin binds the mitochondrial fission GTPase dynamin-related protein-1 and increases its enzymatic activity. Nature medicine. Mar 2011;17(3):377–82. doi:10.1038/nm.2313

56. Panov AV, Gutekunst CA, Leavitt BR, et al. Early mitochondrial calcium defects in Huntington’s disease are a direct effect of polyglutamines. Nature neuroscience. Aug 2002;5(8):731–6. doi:10.1038/nn884

57. Gellerich FN, Gizatullina Z, Nguyen HP, et al. Impaired regulation of brain mitochondria by extramitochondrial Ca2+ in transgenic Huntington disease rats. The Journal of biological chemistry. Nov 7 2008;283(45):30715–24. doi:10.1074/jbc.M709555200

58. Lim D, Fedrizzi L, Tartari M, et al. Calcium homeostasis and mitochondrial dysfunction in striatal neurons of Huntington disease. The Journal of biological chemistry. Feb 29 2008;283(9):5780–9. doi:10.1074/jbc.M704704200

59. Milakovic T, Quintanilla RA, Johnson GV. Mutant huntingtin expression induces mitochondrial calcium handling defects in clonal striatal cells: functional consequences. The Journal of biological chemistry. Nov 17 2006;281(46):34785–95. doi:10.1074/jbc.M603845200

60. Brustovetsky N, Brustovetsky T, Purl KJ, Capano M, Crompton M, Dubinsky JM. Increased susceptibility of striatal mitochondria to calcium-induced permeability transition. J Neurosci. Jun 15 2003;23(12):4858–67.

61. Jędrak P, Krygier M, Tońska K, et al. Mitochondrial DNA levels in Huntington disease leukocytes and dermal fibroblasts. Metab Brain Dis. 08 2017;32(4):1237–1247. doi:10.1007/s11011-017-0026-0

62. Petersen MH, Budtz-Jørgensen E, Sørensen SA, et al. Reduction in mitochondrial DNA copy number in peripheral leukocytes after onset of Huntington’s disease. Mitochondrion. Jul 2014;17:14–21. doi:10.1016/j.mito.2014.05.001

63. Ashraf A, Fan Z, Brooks DJ, Edison P. Cortical hypermetabolism in MCI subjects: a compensatory mechanism? Eur J Nucl Med Mol Imaging. Mar 2015;42(3):447–58. doi:10.1007/s00259-014-2919-z

64. Rudow G, O’Brien R, Savonenko AV, et al. Morphometry of the human substantia nigra in ageing and Parkinson’s disease. Acta Neuropathol. Apr 2008;115(4):461–70. doi:10.1007/s00401-008-0352-8

65. Lunnon K, Keohane A, Pidsley R, et al. Mitochondrial genes are altered in blood early in Alzheimer’s disease. Neurobiol Aging. 05 2017;53:36–47. doi:10.1016/j.neurobiolaging.2016.12.029

66. Mastroeni D, Khdour OM, Delvaux E, et al. Nuclear but not mitochondrial-encoded oxidative phosphorylation genes are altered in aging, mild cognitive impairment, and Alzheimer’s disease. Alzheimers Dement. May 2017;13(5):510–519. doi:10.1016/j.jalz.2016.09.003

