## Supplemental Material for "Mitochondrial and redox modifications in early stages of Huntington’s disease" (from another version of this preprint: version 1, 2022-01-15)

*Supplemental Materials* (Lopes and Ferreira et al.)

### *Supplemental data*

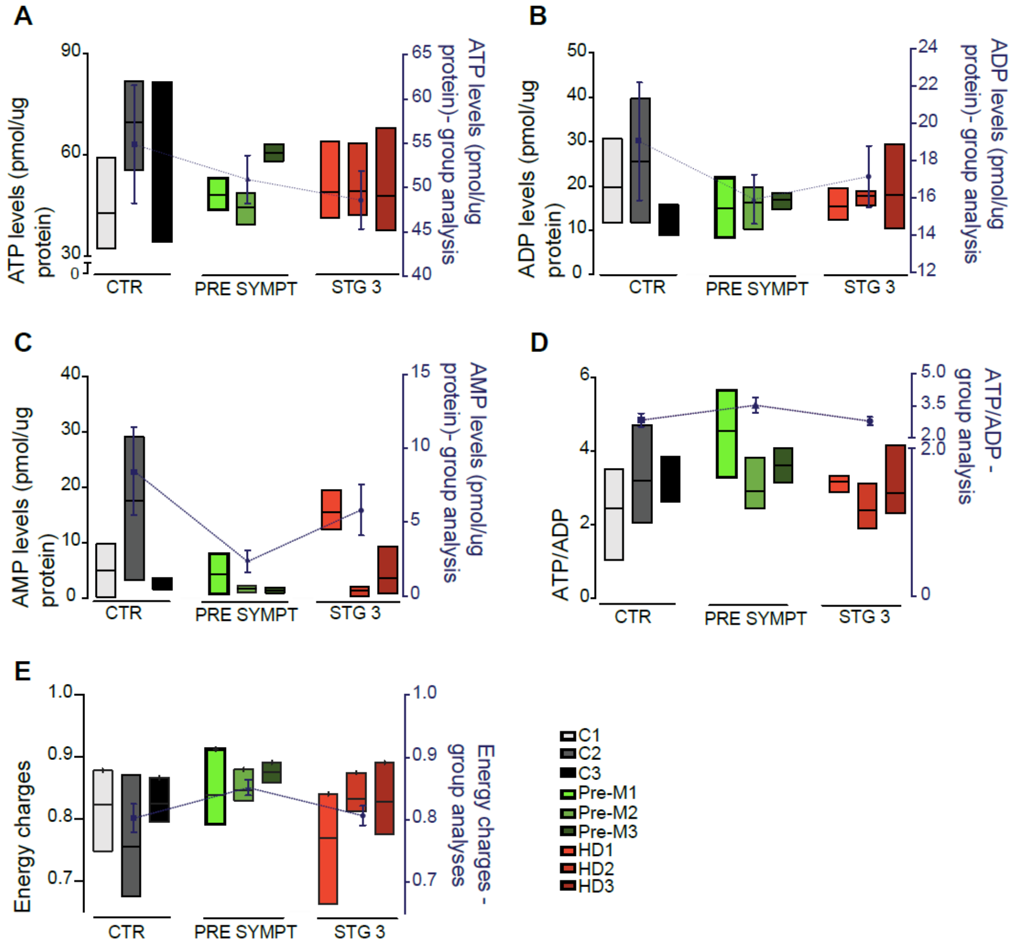


SUPPLEMENTARY FIGURE S1: **Intracellular levels of adenine nucleotides, ATP, ADP and AMP in cultured HD carriers and control human fibroblasts. (A-C)** Cell lysates were assayed for ATP, ADP, and AMP by separation in a reverse-phase HPLC with detection at 254 nm. Individual data are presented as floating bars (min, max) with mean lines shown at least 3 independent experiments; group analysis represents the mean±SEM values (controls (CTR), premanifest (Pre-M) and manifest HD patients) and are plotted on the secondary Y axis (*blue*).

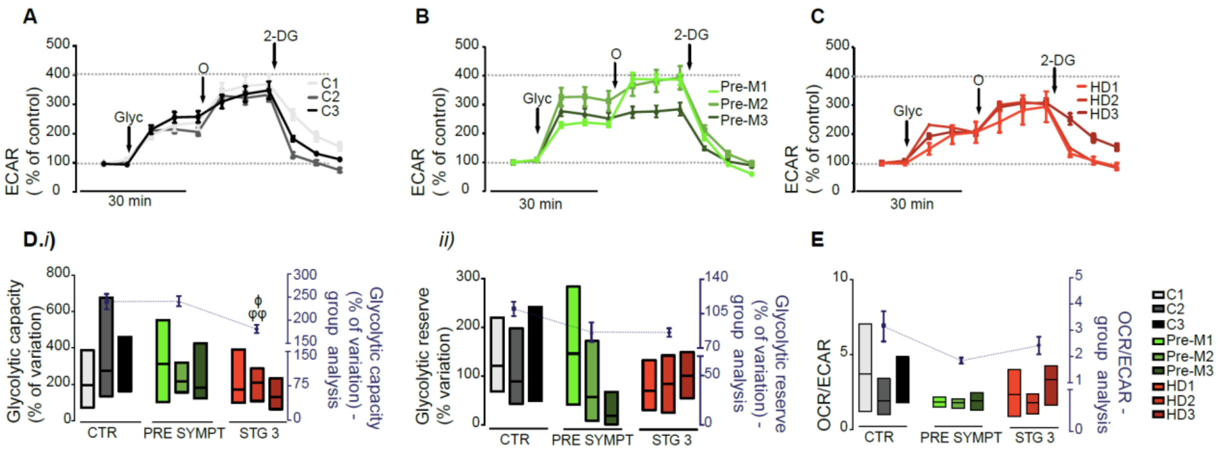

SUPPLEMENTARY FIGURE S2**: Analysis of extracellular acidification rate (ECAR) and OCR/ECAR ratio in skin fibroblasts from HD carriers and controls. (A-C)** ECAR of cells treated with the different compounds. Cells were maintained in assay medium with 0 mM glucose for 1h before the assay. ECAR changes were determined after addition of glucose (Glyc) (10 mM), oligomycin A (O) (1 μM) and 2-deoxy-D-glucose (2-DG) (50 mM) is shown. **(D)** (*i*) Levels of glycolytic capacity and (*ii*) glycolytic reserve. **(E)** OCR/ECAR ratio. Individual data are presented as floating bars (min, max) with mean lines shown at least 3 independent experiments; group analysis represents the mean±SEM values (controls (CTR), premanifest (Pre-M) and HD manifest patients) and are plotted on the secondary Y axis (blue). One-Way ANOVA followed by post hoc Bonferroni's multiple comparisons test. ϕ $p<0.05$ (controls vs HDs), φφ $p<0.01$ (premanifest vs HDs).

SUPPLEMENTARY TABLE S1: **Detailed morphological parameters in mitochondria of HD human fibroblasts and controls.**

| | | Median | Mean (SEM) | Min | Max | Skewness |
|---|---|---|---|---|---|---|
| **Surface area, μm²** | | | | | | |
| | **Ctr** | 0.47 | 0.51 (0.028) | 0.11 | 1.27 | 0.52 |
| | **presympt** | 0.43 | 0.51 (0.041) | 0.07 | 1.86 | 1.72 |
| | **Stg 3** | 0.41 | 0.49 (0.029) | 0.05 | 1.80 | 1.12 |
| **Circularity (0-1)** | | | | | | |
| | **Ctr** | 0.93 | 0.88 (0.01)**** | 0.54 | 1 | −0.85 |
| | **presympt** | 0.71 | 0.73 (0.02)φφφ | 0.29 | 1 | −0.37 |
| | **Stg 3** | 0.88 | 0.82 (0.02)Φ | 0.29 | 0 | −0.79 |
| **Feret's Diameter, μm** | | | | | | |
| | **Ctr** | 0.47 | 0.50 (0.02)* | 0.19 | 1.17 | 1.02 |
| | **presympt** | 0.58 | 0.61 (0.03)φ | 0.12 | 1.59 | 1.05 |
| | **Stg 3** | 0.44 | 0.52 (0.02) | 0.12 | 1.27 | 0.81 |
| **Perimeter, μm** | | | | | | |
| | **Ctr** | 2.69 | 2.68 (0.09) | 1.15 | 5.61 | 0.63 |
| | **presympt** | 2.74 | 0.86 (0.13) | 0.95 | 6.12 | 0.81 |
| | **Stg 3** | 2.49 | 2.73 (0.11) | 0.78 | 5.94 | 0.59 |
| **Form factor** | | | | | | |
| | **Ctr** | 1.06 | 1.24 (0.04)**** | 1 | 2.3 | 1.55 |
| | **presympt** | 1.5 | 1.57 (0.06)φφφ | 1 | 3.95 | 2.08 |
| | **Stg 3** | 1.16 | 1.34 (0.04) | 1 | 3.5 | 2.00 |
| **Aspect ratio** | | | | | | |
| | **Ctr** | 1.41 | 1.78 (0.09)**** | 0.83 | 4.29 | 1.26 |
| | **presympt** | 2.5 | 2.65 (1.72)φφφ | 1.07 | 7.67 | 1.72 |
| | **Stg 3** | 1.66 | 2.06 (1.61) | 0.87 | 6.77 | 1.61 |

Descriptive statistic of mitochondria area, circularity, Feret's diameter, perimeter, form factor and aspect ratio. Statistical analysis: ANOVA followed by pos-doc Bonferroni test: **** p<0.0001 (controls vs premanifest); ϕ p<0.05, (controls vs HDs) and φ p<0.05, φφφ p<0.001 (premanifest vs HDs).

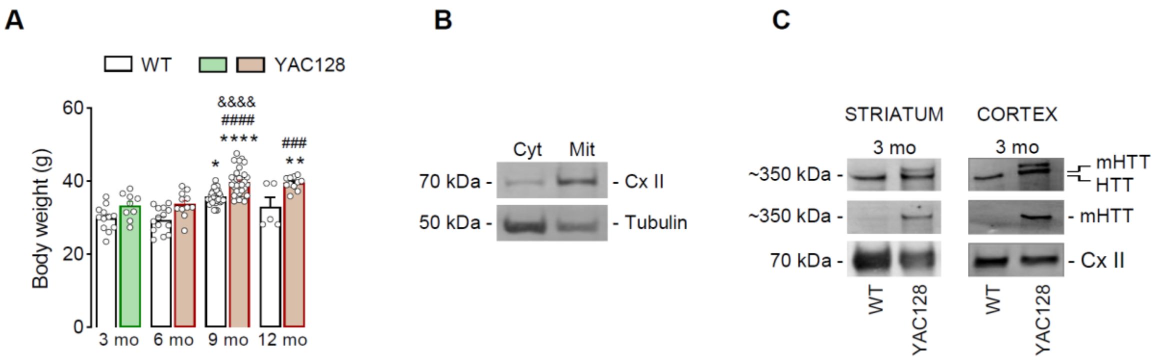


SUPPLEMENTARY FIGURE S3: **Characterization of the YAC128 vs WT mice and mitochondrial fractions.** WT and YAC128 mice body weight at 3, 6, 9 and 12 mo (**A**) and representative western blots indicating the purity grade of mitochondrial-enriched fractions (**B**) stageand the presence of HTT and mHtt in striatal and cortical isolated mitochondria from 3 mo YAC128 mice using anti-HTT antibody (MAB2166) or anti-polyQ antibody (MAB1574-1C2) **(C)**. Data are mean ± SEM of 5-31 mice for body weight analysis. Statistical analysis: *p<0.05; ****p<0.0001 when comparing between genotypes; ####p<0.0001 when compared to 3 mo YAC128 mice; &&&&p<0.0001 when compared to 6 mo YAC128 mice, by two-way ANOVA followed by Tukey's post-hoc test.

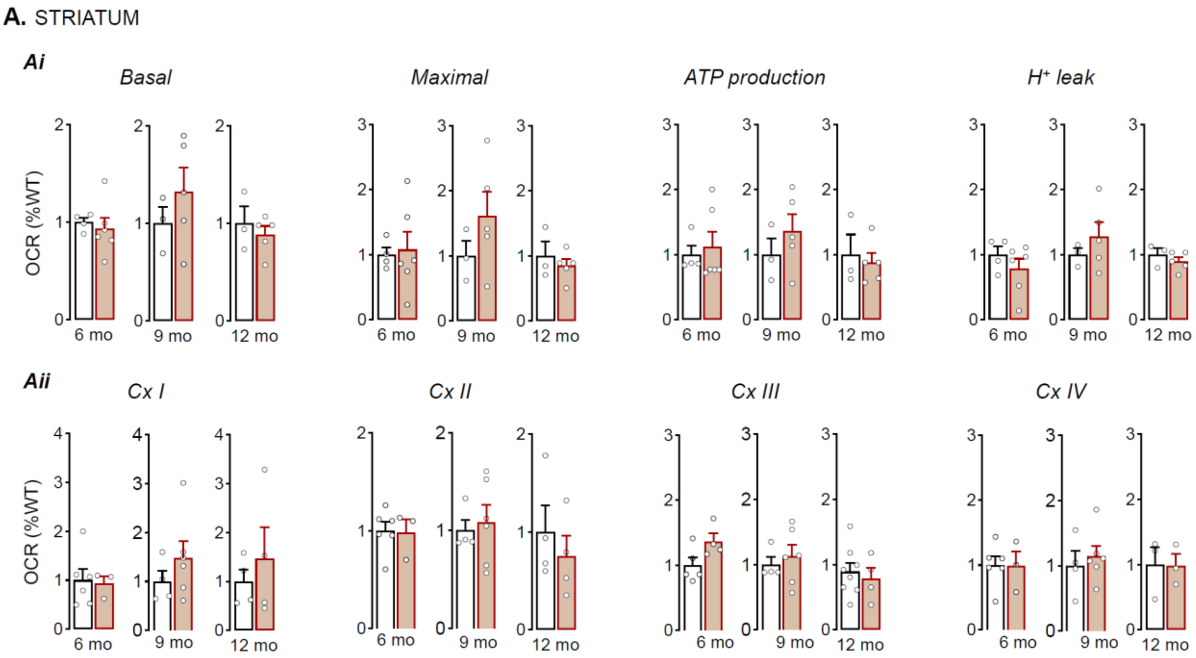

**B.** CORTEX

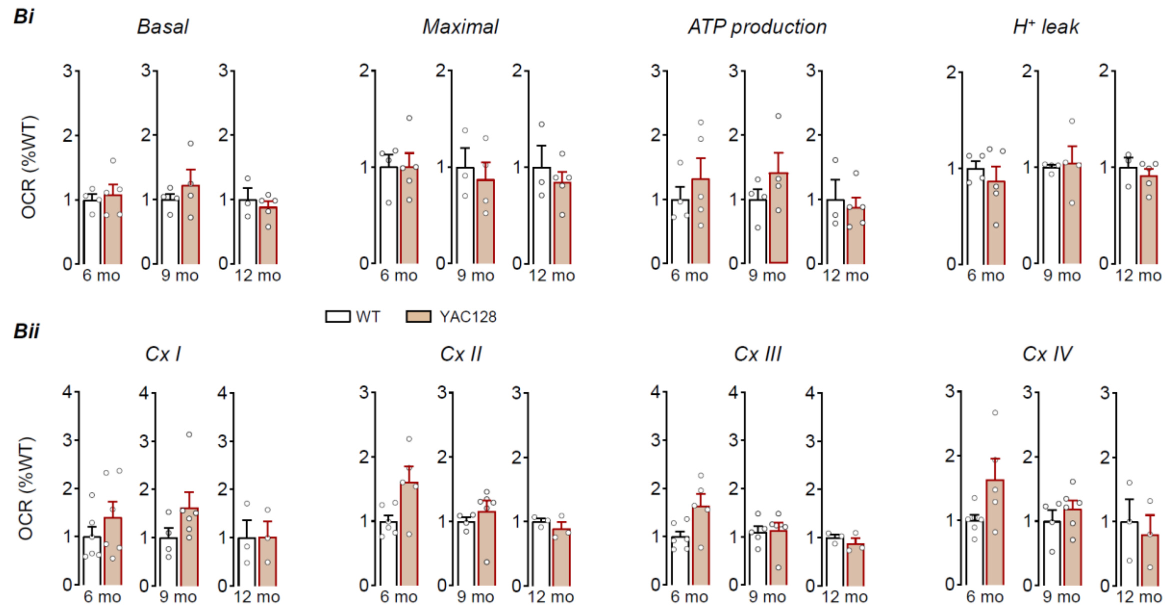


SUPPLEMENTARY FIGURE S4**: Oxygen Consumption Rate (OCR) and respiratory complex activities in sriatal and cortical mitochondria isolated from YAC128 and WT mice at 6-12 months of age.** Levels of respiratory coupling and activity of mitochondrial respiratory chain complexes Cx I-IV was measured in isolated striatal (**A**) and cortical (**B**) mitochondria from 6, 9 and 12 mo YAC128 and WT mice by using the Seahorse XF24 Analyzer. Basal respiration, maximal respiration, ATP production and $H^+$ leak were evaluated in MAS containing 10 mM succinate plus 2 µM rotenone under sequentially injection of mitochondrial inhibitors and substrates (final concentration: 4 mM ADP; 2.5 µg/mL oligomycin; 4 µM FCCP and 4 µM antimycin A) **(A*i* and B*i*)**. **(B)** Activity of mitochondrial respiratory chain complexes Cx I-IV were evaluated in uncoupling conditions performed in MAS containing 4 µM FCCP, 10 mM pyruvate and 2 mM malate. Mitochondrial inhibitors and substrates were sequentially injected (final concentration: 2 µM rotenone, 10 mM succinate, 4 µM antimycin A and 10 mM ascorbate/100 µM TMPD) (**B*i* and B*ii***). Results were calculated as described in Methods section. Data are the mean ± SEM of experiments performed in independent mitochondrial preparations from 3-7 mice from each genotype, run in duplicates or triplicates.

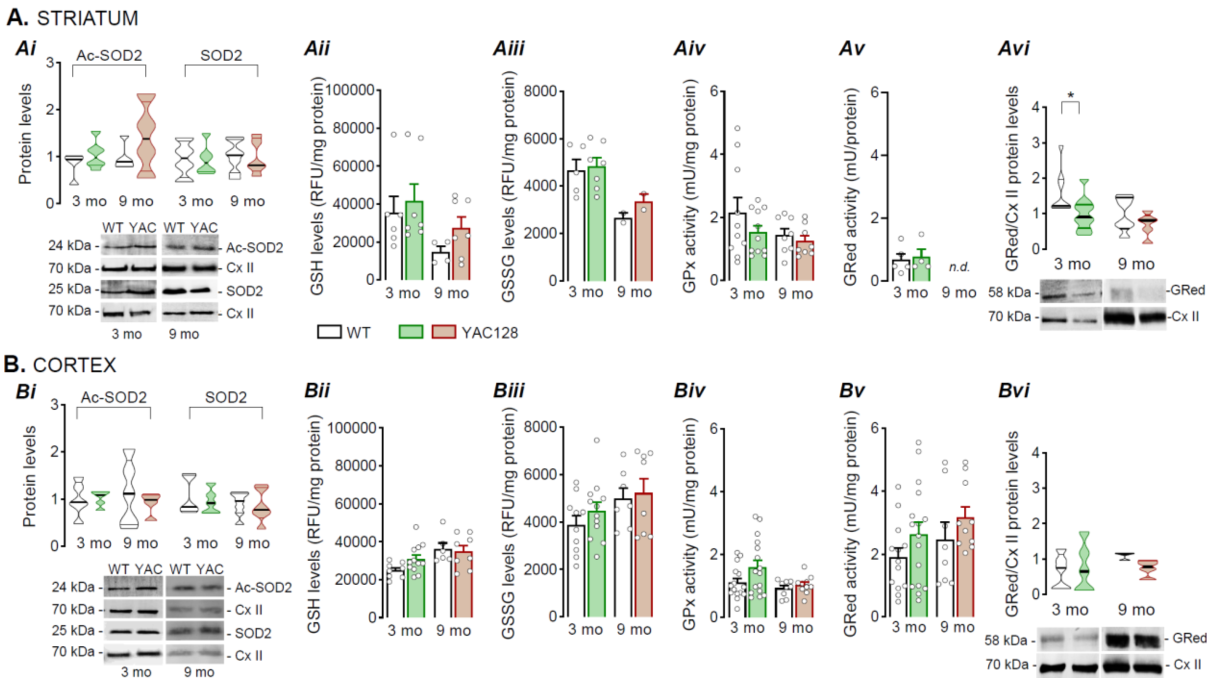

SUPPLEMENTARY FIGURE S5: **Total and acetylated (K68) SOD2 protein levels and glutathione redox cycle evaluation in striatum and cortex of YAC128 and WT mice.** Striatal (A) and cortical (B) mitochondrial extracts from 3 and 9 mo YAC128 and WT mice were evaluated for levels of total acetyl (K68)SOD2 and SOD2 (**A*i* and B*i***) by Western blotting, using 70 kDa subunit of CxII as reference. Total levels of GSH **(A*ii* and B*ii*)** and GSSG **(A*iii* and B*iii*)** and glutathione peroxidase (GPx) **(A*iv* and B*iv*)** and reductase (GRed) **(A*v* and B*v*)** activities were analyzed, as described in Methods section, by spectrophotometric and fluorimetric assays, respectively, in 3 and 9 mo YAC128 and WT mice striatal and cortical isolated mitochondria. Levels of GRed protein levels were performed in mitochondrial extracts obtained from 3 and 9 mo YAC128 and WT mice cortex or striatum by Western blotting, using 70 kDa subunit of Cx II as reference **(A*vi* and B*vi*)**. Data are the mean ± SEM of experiments performed in independent mitochondrial preparations from 2-16 mice from each genotype, run in duplicates for GSH and GSSG levels or GPx and GRed activities. For Western Blotting extracts from 3-11 mice cortex and striatum, from each genotype, were analyzed. Statistical analysis: *$p<0.05$ performed by nonparametric Mann-Whitney U test.

***Supplemental Methods***

**Cell culture**

Fibroblasts were generated from a small skin sample. Briefly, skin explants were washed in culture media several times and cut into small pieces using sterile forceps and scalpel. A few drops of media (well separated from each other) was added onto a 60 mm plate and the minced biopsy pieces placed on to each drop. Carefully a sterile cover slip was placed on top of each piece to hold them in place against the bottom of the plate. A few more drops of media were added into the surroundings of the coverslips to prevent samples from drying out and incubate at 37°C, 5% $CO_2$. On day 2 biopsies were checked and a few more drops of media were added to minimize moisture loss. The procedure was maintained until the outgrowth of primary skin cells was visible. Fibroblasts were maintained at 37 °C in humidified incubator containing 95% air and 5% $CO_2$ and harvested by trypsinization with trypsin-EDTA (0.05%, Thermo Scientific). Cell subculture was performed upon confluence was reached and using a 1:3 split ratio.

**Genotyping**

All animals were genotyped by common procedures using an ear-tip DNA polymerase chain reaction (PCR) standard procedure by using LyA1 (CCT GCT CGC TTC GCT ACT TGG AGC), LyA2 (GTC TTG CGC CTT AAA CCA ACT TGG), RyA1 (CTT GAG ATC GGG CGT TCG ACT CGC), RyA2 (CCG CAC CTG TGG CGC CGG TGA TGC), actin forward (GGA GAC GGG GTC ACC CAC AC), and actin reverse (AGC CTC AGG GCA TCG GAA CC) primers. PCR protocol consisted in 35 cycles, with the following temperatures and times for each step: denaturation at 94°C for 30 s; annealing at 63°C for 30 s; and elongation at 72°C for 30s. PCR products were analyzed on 1.7% agarose gels followed by staining with GreenSafe (NZYtech, Lisboa, Portugal) at 100V for 40 minutes with Tris-acetate-EDTA running buffer and visualized under a UV trans-illuminator, Gel Doc XR system (BioRad®, Hercules, USA).

**Bioenergetic assay**

Fibroblasts were seeded 30 000 cells/per well onto an XF24 or XF96 Cell Culture Microplate and allowed to adhere for 24 h at 37ºC. For OCR and ECAR, 1 ml of XF Calibrant Solution was placed into each well of the sensor hydration microplate and the sensor cartridge placed onto the microplate. The plate was incubated with immersed sensors in a non-$CO_2$ incubator at 37°C for ~16 h (overnight). For OCR analysis, XF assay medium (DMEM) was supplemented with 4.5g/L glucose, 4 mM glutamine and 134 mM NaCl pH adjusted to 7.4, at 37°C; whilst for ECAR no glucose was added. The cell medium was aspirated from each fibroblast–containing microplate without disturbing the cell monolayers attached on the well bottom. Gently, each well was washed with 0.5 ml of XF assay medium at 37 °C. Then, 450 µl of XF assay medium at 37°C was added to each well and the plate placed into a non-$CO_2$ incubator at 37°C for 1 h. OCR was measured after the addition of 4 compounds in successive injections: 1 µM oligomycin; 0.3 µM FCCP; and 1 µM rotenone/antimycin and the ECAR was analysed following the sequential addition of: 10 mM glucose, 1 µM oligomycin and 50 mM 2-deoxyglucose (DOG). Data was normalized using sulforhodamine B[32].

OCR was also measured in fresh cortical and striatal mitochondria isolated from YAC128 *versus* WT mice (5 µg of protein per well), attached to XF24 culture plates pre-coated with polyethyleneimine (PEI, 1:15000 dilution from a 50% solution) in mitochondrial assay solution (MAS) (70 mM sucrose, 220 mM mannitol,10 mM $KH_2PO_4$, 5 mM $MgCl_2$, 2 mM HEPES, 1 mM EGTA and 0.2% (w/v) fatty acid-free BSA, pH 7.2 at 37ºC). For this purpose, the multiwall plate was centrifuged at 2,000 *xg* for 20 minutes at 4°C. In order to evaluate the mitochondria attachment efficiency, the plates were analyzed under light microscopy using 20x magnification to ensure consistent adherence to the wells (data not shown). After an incubation period of 8 minutes at 37°C, the multiwell plate was transferred to the Seahorse XF24 flux analyzer.

Two different experiments were performed: 1) the levels of respiratory coupling ("coupling experiment") and 2) the sequential determination of Cx I-IV-dependent respiration ("electron flow experiment") in YAC128 and WT mice striatal and cortical mitochondria. These experiments were performed by using different substrates, modulators and inhibitors, as described in figure legends and in the results section. For the coupling experiments, basal mitochondrial OCR were analyzed in MAS containing: succinate (10 mM; Cx II substrate) plus rotenone (2 mM; Cx I inhibitor). Mitochondria were then energized by adding ADP (4 mM); respiration drived by ATP synthesis was then prevented by addition of oligomycin (2.5 µg/mL; inhibitor of ATP synthase). The addition of the uncoupler FCCP (4 µM) caused an increase in OCR reflecting the maximal respiratory chain activity as well as the maximal substrate oxidation rate; finally, antimycin A (4 µM; Cx III inhibitor) was added to fully block the respiratory chain and the residual OCR. In order to follow electron flux through the electron transport chain

(ETC) and thus each complex activity, another set of experiments was performed. For this purpose, isolated mitochondria were analyzed in uncoupled state by using MAS containing 4 mM FCCP plus 10 mM pyruvate and 2 mM malate. Next, addition of rotenone led to respiration inhibition; further, succinate injection allowed mitochondria to respire by increasing OCR. Electron flow was then evaluated by adding antimycin A causing OCR inhibition as expected. Finally, injection of ascorbate and TMPD (10 mM and 100 μM, respectively), two artificial substrates used to measure Cx IV activity. Ascorbate helps to maintain TMPD in the reduced form acting as electron donor to cytochrome C/Cx IV, increasing OCR values.

For the respiratory coupling experiments, basal and maximal OCR, ATP production through the Cx V and $H^+$ leak which indicates passive $H^+$ leakage across the MIM, were calculated according to the "rate measurement equations" described in[31].

**Measurement of adenine nucleotides**

For ATP measurement, cells were harvested and washed with ice-cold PBS followed by extraction with 0.6 M perchloric acid. Pellet was dissolved in NaOH and supernatant neutralized with 10M KOH. The resulting supernatants were assayed for ATP determination by separation in a reverse-phase high-performance liquid chromatography (HPLC), with detection at 254 nm, as described previously[66]. The samples were assayed by separation in a reverse-phase HPLC, using a Lichrospher 100 RP-18 (5 µm) HPLC column from Merck (Darmstadt, Germany). Peak identity was determined by following the retention time of standards. Cellular energy charge was calculated using the formula: ([ATP] + 0.5 [ADP])/([ATP] + [ADP] + [AMP]) [67]. Results were normalized to protein content.

**Measurement of cellular hydrogen peroxide production and mitochondrial hydrogen peroxide levels**

Fibroblasts were plated 15 000 cells/per well of 96-well assay plates and cultured for 24 h. Prior to acquisition, fibroblasts were washed with HBSS (in mM: 137.9 NaCl, 1.3 $CaCl_2$, 0.5 $MgCl_2$-6$H_2O$ 0.4 MgSO4-7$H_2O$, 5.3 KCl, 0.4 $KH_2PO_4$, 4.2 $NaHCO_3$, 0.3 $Na_2HPO_4$, 5.6 D-glucose, at pH 7.4). Cells were incubated for 20 minutes with either 10 µM of mitochondria peroxy yellow 1 (MitoPY1) (Sigma) or 50 µM Amplex Red reagent and 0.2 U/mL horseradish peroxidase from the Amplex® Red Catalase Assay Kit (Molecular Probes) at 37ºC and 5% $CO_2$. MitoPY1 is a fluorescent probe that selectively targets the mitochondria and detects fluxes of $H_2O_2$ [68], whereas Amplex measures extracellular $H_2O_2$ production. In the presence of $H_2O_2$ the Amplex® Red reagent produces resorufin, a red-fluorescent oxidation product [69]. MitoPY basal levels were measured for 10-15 minutes, followed by exposure to 3 µM myxothiazol (Sigma)[33] and measured for an additional 30 minutes. MitoPY1 and Amplex fluorescence were acquired using a Microplate Spectrofluorometer Gemini EM (Molecular Devices, USA). MitoPY1 was recorded with an excitation light at 503 nm and emission at 530 nm. Resorufin has absorption and fluorescence emission maxima approximately 571 nm and 585 nm. The results were calculated as RFU per 15 000 cells.

$H_2O_2$ production by YAC128 and WT striatal and cortical mitochondria was measured fluorimetrically by the Amplex® Red method up to 10 minutes, at 37°C, using 571 nm excitation and 585 nm emission (Spectrofluorometer Gemini EM). Briefly, 5 µg mitochondria was resuspended in reaction buffer containing Amplex® Red reagent (10 µM) and horseradish peroxidase (0.5 U/mL), supplemented with 3 mM succinate. Results were expressed in % of mito $H_2O_2$ production related to WT mitochondria.

**Measurement of glutathione levels**

Levels of reduced and oxidized glutathione (GSH and GSSG, respectively) were determined in YAC128 and WT striatal and cortical mitochondria by using a fluorimetric assay, according to [20]. GSH levels were measured in samples (20 µg) after the addition of ortho-phthalaldehyde (OPA; 1 mg/mL methanol) and phosphate buffer (100 mM $NaH_2PO_4$, 5 mM EDTA, pH 8.0). After 15 min of incubation at room temperature in the dark, the end point fluorescence was measured using an excitation wavelength of 350 nm and an emission wavelength of 420 nm. For GSSG determinations, 25 µg of samples were mixed during 5 min with N-ethylmaleimide (NEM; 5 mg/mL methanol) which form adducts with GSH. Then, the mixture was incubated for 15 min in 100 mM NaOH plus OPA (1 mg/mL in methanol), and the fluorescence measured with excitation at 350 nm and emission at 420 nm. Both GSH and GSSG levels were quantified in a Microplate Spectrofluorometer Gemini EM (Molecular Devices, USA). The results were calculated and plotted as RFU per mg protein.

**Measurement of glutathione peroxidase (GPx) and glutathione reductase (GRed) activities**

Glutathione peroxidase (GPx) and glutathione reductase (GRed) activities were measured in YAC128 and WT striatal and cortical isolated mitochondria, spectrophotometrically, at 340 nm, through the kinetic analysis of NADPH oxidation, as described previously [20]. Briefly, the activity of GPx in samples (15 μg) was measured upon a 5 min incubation with phosphate buffer containing 0.25 M $KH_2PO_4$, 0.25 M $K_2HPO4$ and 0.5 mM EDTA, pH 7.0, 10 mM GSH and GRed (1 unit) in the dark at 30°C. After the addition of 2.5 mM NADPH and 12 mM tert-butyl hydroperoxide the absorbance was registered for 5 minutes. For the activity of GRed, samples (20-25 μg) were ressuspended in phosphate buffer containing 0.2 M $K_2HPO_4$ and 2 mM EDTA, pH 7.0, plus 2.5 mM NADPH and activity recorded after the addition of 20 mM GSSG. GRed and GPx activities were determined using a Microplate Spectrophotometer SpectraMax Plus384 (Molecular Devices, USA). Results were presented per mg protein.

**Transmission electron microscopy (TEM)**

Fibroblasts were pelleted by centrifugation at 927 xg for 5 minutes. Both pellet cells and about 1 $mm^3$ of striatal tissue were fixed with 2.5% glutaraldehyde in 0.1 M sodium cacodylate buffer (pH 7.2) for 2h or overnight at 4°C and processed as previously described[33]. Following rinsing in the same buffer, post-fixation was performed using 1% osmium tetroxide for 1h. After rinsing in buffer, buffer and distilled water and a final rinsing step in distilled water, 1% aqueous uranyl acetate was added to the cells during 1 h, for contrast enhancement. After rinsing in distilled water, samples were dehydrated in a graded ethanol series (70–100%). Following embedding in 2% molten agar, cell pellets were re-dehydrated in ethanol (30-100%), impregnated and included in Epoxy resin (Fluka Analytical). Ultrathin sections were mounted on copper grids and stained with lead citrate 0.2%, for 7 min.

Mitochondrial morphometric parameters: Surface area (mitochondrial size) is described in squared micrometers ($\mu m^2$); perimeter in μm; circularity [4(surface area/$perimeter^2$)], indexes of sphericity with values of 1 indicating perfect spheroids; form factor (FF) [($perimeter^2$)/(4·surface area)] reflects the complexity and branching aspect of mitochondria, Feret's diameter represents the longest distance (μm) between any two points within a mitochondrion and aspect ratio (AR) is calculated as [(major axis)/(minor axis)] and reflects the "length-to-width ratio" [70].

All samples were examined with a FEI-Tec-nai G2 Spirit Bio Twin at 100kV at the Bio-Imaging and Electron Microscopy Service at Institute for Biomedical Imaging and Life

Sciences, University of Coimbra. Mitochondrial morphometric parameters were obtained using Image J by manually tracing the longer and smaller mitochondrial axis on TEM micrographs.

**Mitochondrial extracts preparation and Western blotting**

*Ex vivo* striatal- and cortical-enriched mitochondrial fractions were isolated from 3-12-month-old YAC128 and age-matched WT mice. For this purpose, cortical or striatal tissues were homogenized at 4°C in sucrose buffer (in mM): 250 sucrose, 20 HEPES, 100 KCl, 1.5 $MgCl_2$, 1 EGTA, 1 EDTA, pH 7.5/KOH, supplemented with 1 mM DTT, 1 mM PMSF, 100 nM okadaic acid, 5 NaF and 1 µg/mL protease inhibitor cocktail (chymostatin, pepstatin A, leupeptin and antipain, Sigma). Lysates were centrifuged at 560 g for 12 minutes at 4°C and the supernatant collected and again centrifuged at 11900 g for 20 minutes at 4°C. The mitochondrial pellet was resuspended in supplemented TNC buffer (10 mM Tris acetate pH 8, 0.5% Nonidet P40, 5 mM $CaCl_2$) containing 1 µg/mL protease inhibitor cocktail. Protein content in mitochondrial fractions was determined by the Bio-Rad protein assay and samples stored at -80°C until use.

Mitochondrial extracts were denatured with 6x concentrated loading buffer (containing 300 mM Tris-HCl pH 6.8, 12% SDS, 30% glycerol, 600 mM DTT, 0.06% bromophenol blue) at 95°C for 5 min. When isolated mitochondria from cortex were desired for western blotting (for HTT and mHTT labeling and for purity grade analysis), samples were boiled in 1% SDS for 10 min and then denatured with 6x loading buffer as described above. Equivalent amounts of protein (15 µg-50 µg) were separated in 6-15% SDS-PAGE gel electrophoresis (accordingly to the molecular weight of proteins of interest) and electroblotted onto polyvinylidene difluoride (PVDF) membranes. The membranes were further blocked with 5% skim milk or BSA in TBS-Tween: 25 mM Tris-HCl, 150 mM NaCl, pH 7.6,)/0.1% Tween-20 (v/v), during 1 h at room temperature and further incubated overnight at 4°C with gentle agitation with primary antibodies prepared in 1% BSA-containing TBS-Tween: SOD2 (acetyl K68; 1:500; Abcam ab 137037;Abcam, Cambridge, UK); Superoxide dismutase 2 (SOD2; 1:5000; Abcam ab13533; Abcam, Cambridge, UK); glutathione reductase (1:2000; Abcam ab16801; Abcam, Cambridge, UK); Cx II (70 kDa subunit; 1:10000; Molecular Probes A11142; Molecular Probes-Invitrogen,Eugene, OR, USA) was used as mitochondrial marker and tubulin (1:1000, Sigma T-6199; Sigma, St. Louis, MO, USA) as cytosolic marker for the mitochondrial fractions. The presence of HTT and mHtt in isolated striatal and cortical mitochondria from 3 mo YAC128 mice was evaluated using anti-HTT antibody (MAB2166; Merck Millipore, Darmstadt, Germany) or anti-polyQ antibody (MAB1574-1C2; Merck Millipore, Darmstadt, Germany). The purity grade of mitochondrial fractions, obtained by subcellular fractionation, was evaluated by western blotting by using anti-Cx II as mitochondrial marker and tubulin as cytosolic markers. Membranes were incubated with anti-mouse or anti-rabbit IgG secondary antibodies prepared in 1% BSA-containing TBS-Tween for 1 h, at room temperature. In order to normalize the amount of protein per lane anti-Cx II was used as loading control for mitochondrial. Immunoreactive bands were visualized by alkaline phosphatase activity after incubation with ECF reagent on BioRad ChemiDoc Touch Imaging System (BioRad, Hercules, USA) and quantified using Image Lab analysis software (BioRad, Hercules, USA).
